# Synergistic regulation of Notch signaling by different O-glycans promotes hematopoiesis

**DOI:** 10.1101/2022.03.21.485194

**Authors:** Ankit Tanwar, Pamela Stanley

**Author notes:** Corresponding Author: Pamela Stanley, Department of Cell Biology, Albert Einstein College of Medicine, New York, NY, 10461 USA.

## Abstract

Glycosylation of Notch receptors by O-fucose glycans regulates Notch ligand binding and Notch signaling during hematopoiesis. However, roles in hematopoiesis for other O-glycans that modify Notch receptors have not been determined. Here we show that the EGF domain-specific GlcNAc transferase EOGT is required in mice for the optimal production of lymphoid and myeloid cells. The phenotype of *Eogt* null mice was largely cell-autonomous, and Notch target gene expression was reduced in T cell progenitors. Moreover, EOGT supported residual Notch signaling following conditional deletion of *Pofut1* in hematopoietic stem cells (HSC). *Eogt:Pofut1* double mutant HSC had more severe defects in bone marrow, and in T and B cell development in thymus and spleen, compared to deletion of *Pofut1* alone. The combined results show that EOGT and O-GlcNAc glycans are required for optimal hematopoiesis and T and B cell development, and that they act synergistically with POFUT1 and O-fucose glycans to promote Notch signaling in lymphoid and myeloid differentiation.

**Key points:**

- O-GlcNAc glycans and EOGT promote lymphopoiesis and myelopoiesis
- EOGT supports Notch signaling in the absence of POFUT1 and O-fucose glycans

## Introduction

Notch signaling is highly conserved and plays crucial roles in cell fate determination and tissue development ^1^. There are four Notch receptors (NOTCH1 to NOTCH4) that can be activated by canonical Notch ligands (DLL1, DLL3, DLL4, JAG1 and JAG2) to induce Notch signaling in mammals. Notch ligand binding and Notch signaling are regulated by glycosylation of the extracellular domain (ECD) of Notch receptors ^2,3^. Structural studies reveal direct interactions between Notch ligands and O-fucose in specific epidermal growth factor-like (EGF) repeats of NOTCH1 ^4,5^. Following Notch ligand engagement, an ADAM metalloprotease cleaves NECD, followed by a second cleavage by the γ-secretase complex. Notch intracellular domain (NICD) complexes in the nucleus with the transcriptional repressor Rbp-Jk, and co-activators including Mastermind-like-1 (MAML1), to induce the expression of Notch target genes, including Hairy enhancer-of-split (*Hes*) and Hairy-related gene families, which regulate the expression of many other genes ^6,7^.

Within the hematopoietic system, Notch signaling plays important roles in regulating different stages of lymphoid and myeloid development ^8,9^. DLL4-induced NOTCH1 signaling is indispensable for T cell development in the thymus. Thus, conditional inactivation of *Notch1* or *Dll4* using *Mx1-Cre* is sufficient to block T cell development ^10,11^. However, *Notch2* is also required for optimal development of early T cell progenitors ^12^. DLL1-induced NOTCH2 signaling is essential for the generation of marginal zone B cells (MZ-B) in the spleen ^13,14^. O-fucose glycans extended by LFNG and MFNG promote the formation of MZ-B cells ^15^.

Consensus sites within EGF repeats in the NECD of Notch receptors carry O-fucose, O-glucose and O-GlcNAc glycans ^16^ (Figure 1). EGF repeats with appropriate consensus sites occur in ∼100 proteins of the proteome, including Notch receptors and Notch ligands ^17,18^. O-fucose is transferred by protein O-fucosyltransferase 1 (POFUT1), which is further extended by the Fringe family of glycosyltransferases (LFNG, MFNG and RFNG). Misexpression of *Lfng* in thymus disrupts T cell development ^19-21^. Conditional deletion of *Pofut1* in the bone marrow leads to the disruption of hematopoiesis with an increase in granulocyte-monocyte progenitors (GMP), and a reduction in common myeloid progenitors (CMP) ^22^. This causes a block in T cell production in thymus and MZ-B cell production in spleen, accompanied by an increase in granulocytes in spleen. Notch ligand binding and Notch signaling are markedly reduced in hematopoietic stem cells (HSC) following conditional deletion of *Pofut1*. However, *Pofut1* deletion by *Mx1-Cre* gives a milder reduction in thymic T cells than deletion of RBP-Jk ^23^, whereas global deletion of *Pofut1* ^24,25^ or RBP-Jk ^26^ causes similarly severe Notch signaling defects, and embryonic lethality. Thus, loss of O-fucose glycans in HSC does not fully abrogate Notch signaling, suggesting that other O-glycans may support Notch signaling in the absence of POFUT1. Synergism between O-fucose and O-glucose glycans on Notch has been observed in *Drosophila* ^27^, and in mammalian cells ^28^. However, in both cases, the effects on Notch signaling appear to be due to reduced expression of Notch receptors lacking both O-fucose and O-glucose glycans at the cell surface. By contrast, the absence of O-fucose glycans from Notch receptors in CHO and embryonic stem cells only slightly reduces NOTCH1 cell surface levels ^29,30^.

**Figure 1.**
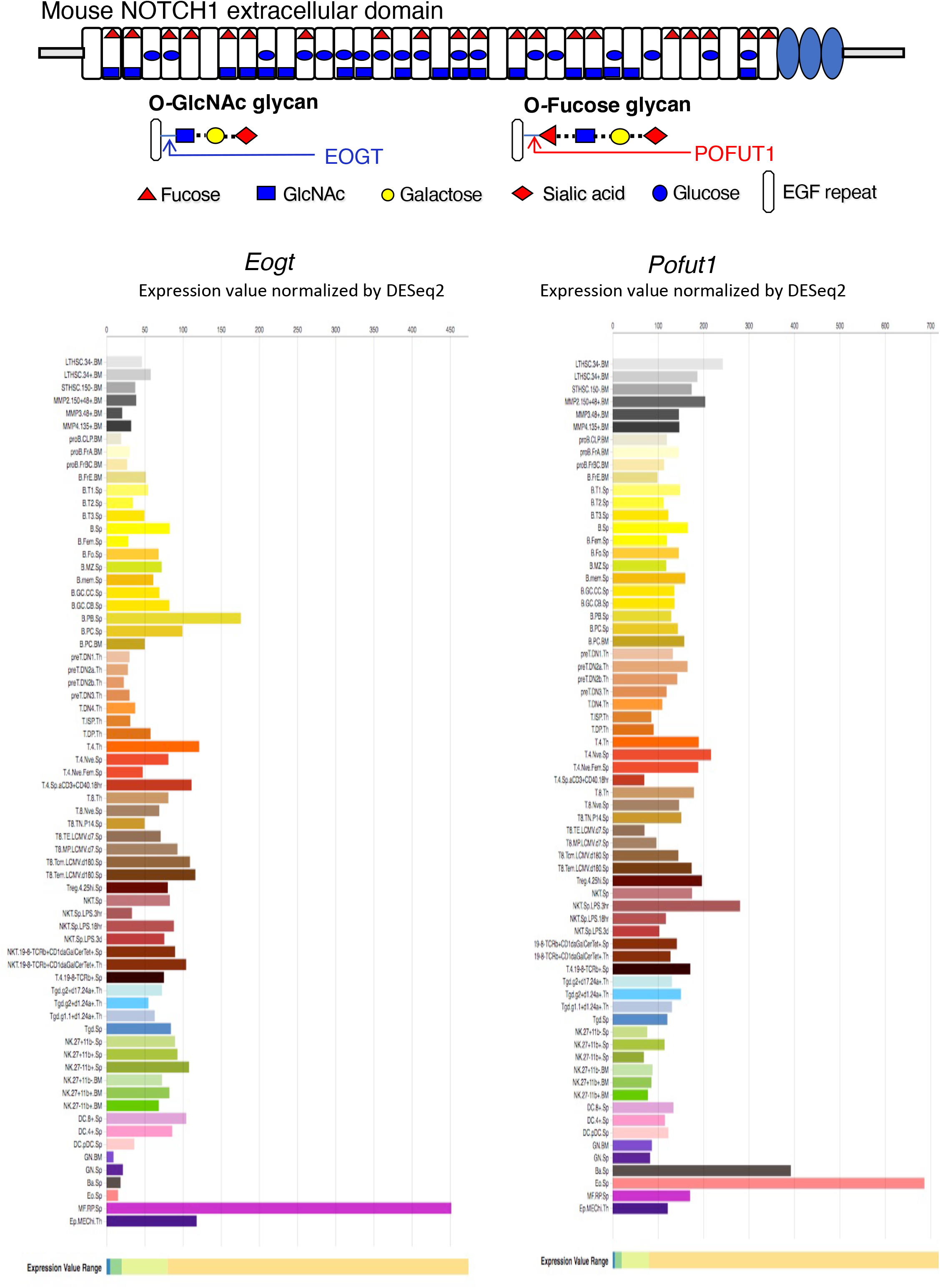
Expression of *Eogt* and *Pofut1* in immune cell subsets. The diagram depicts the ECD of mouse NOTCH1 showing the O-glycans attached at respective consensus sites in the 36 EGF repeats. The sugar that initiates each O-glycan is shown. The potential extensions of O-fucose and O-GlcNAc glycans are also shown. The expression of *Eogt* and *Pofut1* in immune cells is taken from ImmGen Gene Skyline ULI RNASeq data group for *Eogt* and *Pofut1* gene expression ^46^ (http://rstats.immgen.org/Skyline/skyline.html).

The addition of O-GlcNAc to EGF repeats was first identified in *Drosophila* ^31^, and the EGF domain-specific O-GlcNAc transferase EOGT was subsequently identified ^32,33^. *Eogt* null mice exhibit defective postnatal retinal angiogenesis, similar to that observed in mice with disrupted Notch signaling ^34^. Cell-based experiments showed that EOGT promotes the binding of DLL Notch ligands and Notch signaling ^34^. *Eogt* is expressed with *Pofut1* in many cells of the immune system (Figure 1). However, their expression levels vary inversely in some cell types, indicating potentially different functional roles (Figure 1). In this paper, we identify roles for *Eogt* in the regulation of Notch signaling during hematopoiesis, lymphopoiesis and myelopoiesis. In addition, we show that *Eogt* supports Notch signaling in the absence of *Pofut1* and O-fucose glycans.

## Methods

### Mice

Mice with an inactivating mutation in the *Eogt* gene were generated at Nagoya University and previously described ^34^. *Pofut1*[F/F] mice were also previously described ^24^. Transgenic mice expressing *Vav1-iCre* (B6.Cg-*Commd10*^*Tg(Vav1-icre)A2Kio*^/J Strain) were a kind gift from Britta Will and Paul Frenette at the Albert Einstein College of Medicine, NY, USA. Compound mutant mice termed Pof cKO and EPof dKO, with and without *Vav1-iCre*, were generated by intercrossing. C57Bl/6J male congenic mice expressing CD45.1 (*B6*.*SJL-Ptprc*^*a*^ *Pepc*^*b*^/BoyJ *#002014*) were obtained from the Jackson Laboratory (Bar Harbor). Genotyping was performed by PCR of genomic DNA using primers that distinguish wild-type and mutant alleles (supplemental Table1). Mice were housed in a barrier facility, allowed to eat and drink ad libitum, and used in experiments at 7–8 weeks of age. All experiments were performed with permission from the Albert Einstein Institutional Animal Care and Use Committee under approved protocol numbers 20170709 and 00001311. Euthanized mice were weighed, and isolated bone marrow (femurs, tibias, fibulae, & patullae), thymus and spleen were also weighed before making single-cell suspensions.

### Antibodies

Supplemental Table 2 has the full description and commercial source of each antibody used in this work.

### Flow cytometry

Single-cell suspensions were prepared from bone marrow by crushing femurs, tibias, fibulae, and patullae in a mortar and rinsing vigorously with 20 ml cold FACS binding buffer (FBB: Hank’s balanced salt solution (HBSS) with glucose (Corning), 1 mM CaCl_2,_ 2% bovine serum albumin (BSA, fraction V, Sigma) and 0.05% sodium azide, (pH 7.2–7.4). The BM cell suspension was passed through a 70-μm strainer. Thymus or spleen was squeezed through a 70 μm strainer in 1 ml FBB. Thymocytes were washed in 10 ml cold FBB twice. Bone marrow and splenocytes were centrifuged and incubated in 1 ml RBC lysis buffer (eBiosciences) for 3-5 min before adding 10 ml cold FBB. After centrifugation and resuspension in 5 ml cold FBB, single cell suspensions were counted in a Coulter counter. Cells were centrifuged at 4ºC, resuspended in 2 ml cold PBS with 1 mM CaCl_2_ and 1 mM MgCl_2_, pH 7.2, centrifuged and resuspended in 100 μl Zombie NIR dye for live/dead assessment, according to the manufacturer’s protocol (Zombie NIR Fixable Viability Kit, BioLegend). After 30 min at 4ºC in the dark, 2 ml cold FBB was added. Cells were centrifuged at 4ºC and resuspended in 4% paraformaldehyde (PFA, Emsdiasum) in PBS pH 7.2, incubated 15 min at 4ºC in the dark. Cells were washed twice with 2 ml cold FBB, resuspended at 10^6^ cells/ml in FBB and stored at 4ºC for up to 3 months. For analysis by flow cytometry, ∼10^6^ cells were washed with 1 ml FBB, resuspended in 90 μl FBB containing 1 μl Fc block (rat-anti-mouse CD16/CD32), and incubated for 15 min on ice. Ab diluted in FBB (10 μl) was added and the reaction mix was incubated for 30 min at 4°C. Cells were washed twice in 1 ml FBB and resuspended in ∼500 μl FBB. For all samples, immunofluorescence was analyzed using a Cytek™ Aurora and BD LSRII flow cytometer and data FCS files were analyzed using FlowJo software (Tree Star). Gating strategies shown in supplemental Figures 1-3 and Figure 6 were based on previous work ^35-37^.

### Bone marrow transplantation

Cell suspensions from bone marrow of 7–8 week *Eogt*[+/-] and *Eogt*[-/-] males were prepared as described above. Approximately 3×10^6^ cells were resuspended in 50 μl sterile HBSS (Gibco) and injected using a 28-gauge insulin needle via the retro-orbital plexus into CD45.1+ C57BL/6 lethally-irradiated recipients. A split dose of 550 rads γ-irradiation per recipient male was given twice, with a 16 h interval. After 7 weeks, recipients were euthanized, bone marrow, thymus, and spleen were analyzed for lymphoid and myeloid cell subsets by flow cytometry after gating on donor-derived cells positive for anti-CD45.2-Pacific Blue.

### Isolation of CD4/CD8 DN T cells

Fresh thymocytes were resuspended in isolation buffer (PBS lacking cations, pH 7.2-7.4, containing 0.1% BSA, 2 mM EDTA and 1 mg/ml glucose) on ice. For T cell depletion, ∼3-5×107 thymocytes were incubated with 20 μg anti-CD4 (rat IgG2b clone GK1.5, BioXCell) and 37.5 μg anti-CD8a (rat IgG2a clone 53-6.72; BioXCell) in 5 ml buffer for 20 min at 4°C with tilted rotation. After centrifugation, cells were resuspended in 5 ml buffer, and incubated with 250 μl sheep anti-rat IgG Dynabeads (Thermo Fisher Scientific) for 30 min at 4°C with tilted rotation. The tube was placed in a magnet for 2 min, unbound cells were centrifuged and resuspended in 250 μl Dynabeads for a second 30 min incubation at 4°C. After Dynabeads removal, unbound cells were centrifuged, counted and RNA was extracted from the cell pellet with 1 ml TRIZOL (Ambion) as described below.

### Quantitative RT-PCR

DN T cells (∼3×10^7^) thymocytes were pipetted vigorously in 1 ml TRIZOL and RNA was isolated according to the manufacturer’s instruction. cDNA was prepared from 250 ng RNA using the ReverTra Ace® qPCR RT Master Mix with gDNA Remover (Dc. DiagnoCine) following the manufacturer’s protocol. Primer sequences used for qRT-PCR are in supplemental Table1.

### Histopathology

Spleen was collected, washed and stored in 10% natural buffered formalin (NBF) at 4ºC. The samples were processed for paraffin embedding and longitudinal tissue sections (5 μm) were stained with hematoxylin–eosin (H & E), scanned by the 3D Histech P250 High-Capacity Slide Scanner and analyzed using Case Viewer 2.4 software.

### Notch ligand binding assay

Soluble Notch ligands DLL1-Fc (#10184-DL-050), JAG1-Fc (#1277-JG-050), and JAG2-Fc (#1726-JG-050) were purchased from R&D Systems, and DLL4-Fc (#DL4-H5259) was purchased from Acro-biosystems. Single-cell suspensions from thymus were washed in FBB at 4ºC and fixed in PBS-buffered 4% PFA for 15 min at 4ºC, washed twice with FBB and stored in FBB at 4°C. For analysis, ∼0.5-1×10^6^ fixed thymocytes were washed with FBB, and incubated with 40 μl FBB and 1 μl FcR blocking solution (rat-anti-mouse CD16/CD32) on ice for 15 min. Thereafter, the cells were incubated in 60 μl FBB containing anti-CD4-FITC (1:400), anti-CD8a-PerCp-Cy5.5 (1:400), and 0.75 μg DLL4-Fc, or 1.5 μg of DLL1-Fc, JAG1-Fc, or JAG2-Fc. After incubation at 4°C for ∼30 min, cells were washed with 1 ml FBB and incubated with anti-IgG-APC and anti-IgG-DyLight 405 (Fc-specific) Ab (1:100) at 4°C for 30 min. The cells were then washed with 1 ml FBB and resuspend in 500 μl FBB and analyzed in a flow cytometer (Cytek™, Aurora). For detection of NOTCH1 at the cell surface, fixed DN T cells were incubated with FcR-block rat-anti-mouse CD16/CD32 (1:100) followed by anti-CD4-FITC mAb (1:400), anti-CD8a-PerCp-Cy5.5 (1:400), sheep anti-mouse NOTCH1 Ab (1:50) at 4°C for 30 min, washed, and incubated with rhodamine Red-X-conjugated donkey anti-sheep IgG (1:100) at 4°C for 30 min. Cells were washed with 1 ml FBB, resuspended in 500 μl FBB and analyzed in a FACS flow cytometer (Cytek™, Aurora).

### Statistics

Comparisons are presented as mean ± SEM. Significance was determined by both two-tailed and one-tailed unpaired (denoted by parentheses), parametric, Student t-test analysis (unless otherwise noted) using Prism software version 9.1.

## Results

### Loss of *Eogt* affects myelopoiesis and lymphopoiesis

Initial comparisons of *Eogt*[+/+] and *Eogt*[+/-] heterozygotes revealed no significant differences in T, B and myeloid subsets (supplemental Figure 4). Therefore, data from *Eogt*[+/+] and *Eogt*[+/-] mice were combined as Control. Thymus and spleen weights, as well as bone marrow (BM) cellularity, were similar in *Eogt* null and Control mice (supplemental Figure 5A). In bone marrow, the number of CD19+/B220+ B cells, and CD11b+/Gr1+ granulocytes were significantly increased in the *Eogt*[-/-] population (Figure 2A). In thymus, the frequency of CD4/CD8 double negative 1 (DN1), and the absolute cell number of DN2 T cell progenitors, were strikingly reduced in *Eogt* null cells, while the number of DN4 T cells was significantly increased (Figure 2B). By contrast, the numbers of early T cell progenitors (ETP) and DN3 T cells were unchanged (supplemental Figure 5B). The proportion of double positive (DP) T cells was slightly decreased, while CD4+ and CD8+ single positive (SP) T cells were increased in *Eogt*[-/-] thymocytes (Figure 2B). These effects were also observed in mice lacking the three Fringe genes (*Fng* tKO) ^36,38^. Consistent with inhibition of Notch signaling in the *Eogt* null thymus, there was a significant increase in thymic B cells (CD19+/B220+) and the frequency of myeloid cells (CD11b+) (Figure 2B), but no change in natural killer (NK) T cells (supplemental Figure 5B). In spleen, no histopathological changes were observed in Control versus *Eogt*[-/-] sections (n=3 for each, not shown), and the absolute numbers of T cells, B cells, marginal zone progenitors (MZ-P) and myeloid cell subsets were similar in Control versus *Eogt*[-/-] mice (supplemental Figure 5C). However, significant increases were observed in the number of follicular B (Fo-B), MZ-B, CD19+ and B220+ B cells in *Eogt*[-/-] mice (Figure 2C). By contrast, a decrease in the proportion of natural killer T cells (NK1.1+) and dendritic cells (CD11b/c+) was observed (Figure 2C). Thus, EOGT and O-GlcNAc glycans are required for the optimal generation of lymphoid and myeloid cells from HSC.

**Figure 2.**
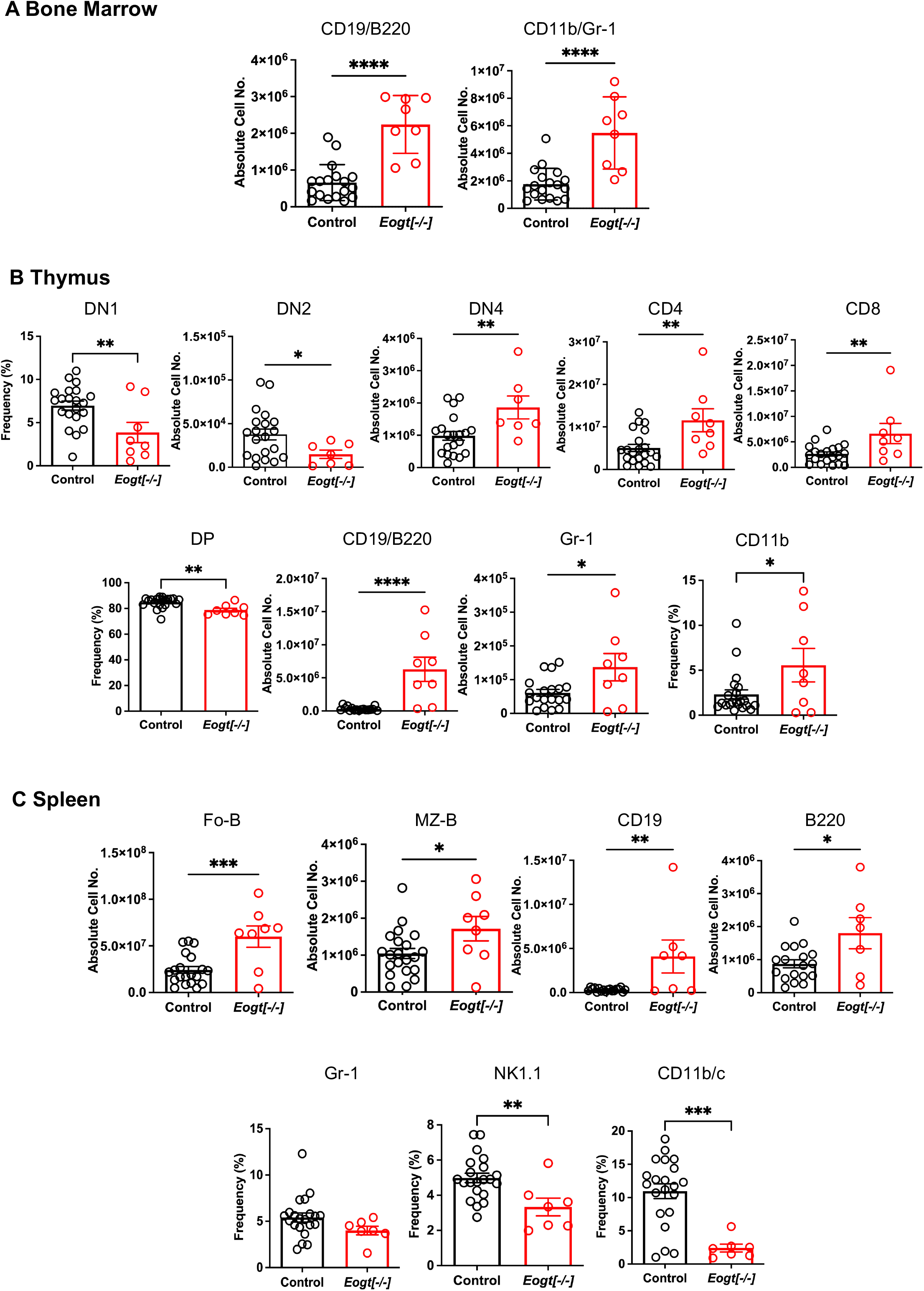
Altered lymphoid and myeloid subsets in mice lacking *Eogt* and O-GlcNAc glycans. (A) Absolute cell numbers or frequency (%) for lymphoid and myeloid cell subsets in bone marrow that differed between Control (*Eogt*[+/+] and *Eogt*[+/-]) and *Eogt*[-/-] mice. See supplemental Figure 1 for gating, supplemental Figure 4 for *Eogt*[+/+] versus *Eogt*[+/-] data and supplemental Figure 5 for BM cellularity. (B) Absolute cell numbers or frequency (%) for lymphoid and myeloid cell subsets in thymus that differed between Control and *Eogt*[-/-] mice. See supplemental Figure 2 for gating, supplemental Figure 4 for *Eogt*[+/+] versus *Eogt*[+/-] data and supplemental Figure 5 for subsets that did not differ significantly. (C) Absolute cell numbers or frequency (%) for lymphoid and myeloid cell subsets in spleen that differed between Control and *Eogt*[-/-] mice. See supplemental Figure 3 for gating, supplemental Figure 4 for *Eogt*[+/+] versus *Eogt*[+/-] data and supplemental Figure 5 for subsets that did not differ significantly. Each symbol represents a mouse of 7-8 weeks. Data are presented as mean ± SEM. *p <0.05, **p<0.01,***p<0.001, ****p<0.0001 based on two-tailed Student t test or (*) p <0.05 based on one-tailed Student t test.

### The *Eogt* null phenotype is largely cell autonomous

To determine whether the *Eogt* null phenotype was cell intrinsic, bone marrow (BM) transplantation was performed from CD45.2+ *Eogt*[+/-] and *Eogt*[-/-] donor males into CD45.1+ male hosts. *Eogt*[-/-] CD45.2+ donor BM cells reconstituted CD45.1+ hosts to ∼59% in bone marrow (Figure 3A), ∼80% in thymus and ∼64% in spleen (not shown). Thus, host-derived *Eogt*[+/+] cells contributed to each of these populations in recipients. In recipient bone marrow, no changes were seen in B cells (CD19+/B220+) or granulocytes (CD11b+/Gr-1+) from *Eogt*[-/-] donor BM compared to *Eogt*[+/-] BM (Figure 3B); in recipient thymus, *Eogt*[-/-] donor BM generated significantly fewer DN2 and DN3 T cell progenitors compared to *Eogt*[+/-] donor BM (Figure 3C); and in spleen, *Eogt*[-/-] donor BM generated CD19+ cells and Gr-1+ granulocytes in significantly increased numbers compared to *Eogt*[+/-] donor BM (Figure 3D). Thus, the phenotype of *Eogt*[-/-] BM recipients was somewhat milder, than the *Eogt* null phenotype (Figure 3B). This could reflect the presence of *Eogt*[+/+] host cells, and/or rescuing effects of the wild type host stroma. Nevertheless, key aspects of the *Eogt* null phenotype were transferred by *Eogt*[-/-] BM.

**Figure 3.**
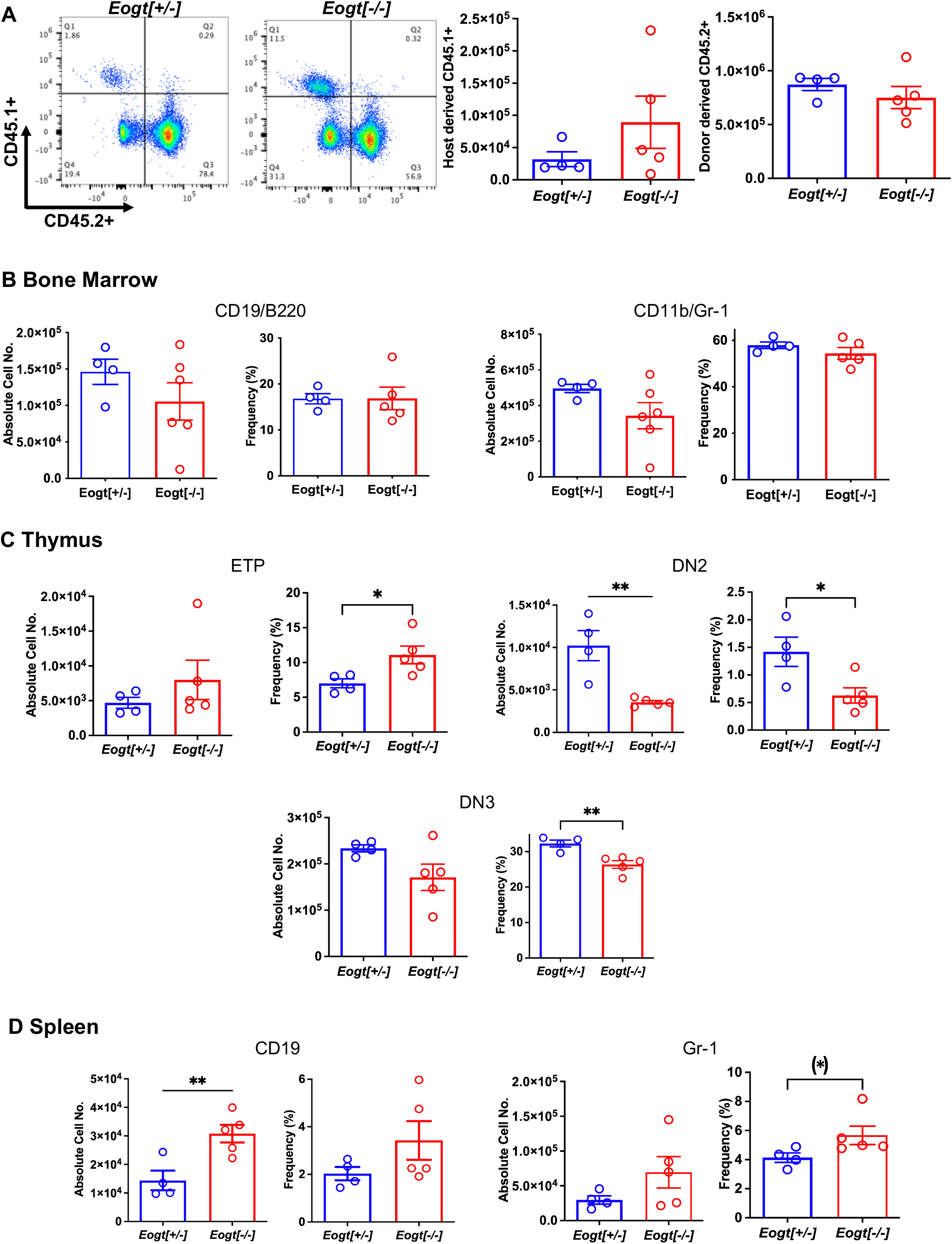
The *Eogt* null phenotype is largely cell autonomous. Bone marrow cells (3×10^6^) from *Eogt*[+/-] or *Eogt*[-/-] mice expressing CD45.2 were injected via the retro-orbital plexus into lethally-irradiated, wild-type recipient males expressing CD45.1. Recipient BM, thymus and spleen were analyzed 7 weeks after transplantation. (A) Flow cytometry profile and histograms of bone marrow from recipient mice (CD45.1+) that received BM from *Eogt*[+/-] or *Eogt*[-/-] mice. The approximate donor contribution was 78% for *Eogt*[+/-] and 57% for *Eogt*[-/-] of recipient BM. (B) Absolute cell number and frequency (%) of B cells and granulocytes in bone marrow that differed when transferred from *Eogt*[+/-] versus *Eogt*[-/-] donor bone marrow. (C) Absolute cell number and frequency (%) of T cell progenitors in thymus that differed when derived from *Eogt*[+/-] versus *Eogt*[-/-] donor bone marrow. (D) Absolute cell number and frequency (%) of B cells and granulocyte subsets in spleen that differed when derived from *Eogt*[+/-] versus *Eogt*[-/-] donor bone marrow. Each symbol represents a mouse of 7-8 weeks. Data are presented as mean ± SEM. *p <0.05, **p<0.01 based on two-tailed Student t test. (*) p <0.05 based on one-tailed Student t test.

### Notch signaling is reduced in *Eogt*[-/-] DN T cell progenitors

Notch ligand binding was examined using thymic DN T cell progenitors from 7-8 week *Eogt*[+/-] and *Eogt*[-/-] mice. No significant changes were observed in either NOTCH1 cell surface expression, or the binding of soluble ligands DLL1, DLL4, JAG1 or JAG2 between Eogt[+/-] and *Eogt*[-/-] DN T cell progenitors (Figure 4A). However, there were significant reductions in the expression of two Notch target genes, *Hes1* and *Il2ra*, consistent with reduced Notch signaling (Figure 4B). The expression of *Il2ra* was also reduced in DN T cell progenitors from mice lacking the three Fringe genes, along with *Dtx1*, although not *Hes1* ^36^. The reduction in Notch signaling target gene expression in *Eogt* null T cell progenitors, the changes in T cell subset numbers and frequencies, and the increased numbers of B and myeloid cells observed in *Eogt*[-/-] thymus, indicate that EOGT and O-GlcNAc glycans are necessary for optimal Notch signaling and T cell development. The altered B cell and myeloid subsets in spleen of *Eogt*[-/-] mice are also consistent with reduced Notch signaling. Finally, the B cell and myeloid hyperplasia in *Eogt*[-/-] BM indicate that Notch signaling is required for regulating the differentiation of certain progenitors during hematopoiesis.

### *Eogt* supports lymphoid and myeloid development in HSC lacking Pofut1

Previous work showed that inactivation of *Pofut1* using *Mx1-Cre* causes a reduction in T lymphopoiesis in thymus and myeloid hyperplasia in bone marrow ^22^. In another study, conditional deletion of *Pofut1* in bone marrow was shown to cause a milder T cell phenotype than deletion of RBP-Jk ^23^. To determine whether *Eogt* and O-GlcNAc glycans support Notch signaling and hematopoiesis in the absence of *Pofut1*, we used *Vav1-iCre* transgenic mice to generate conditional deletion (cKO) of *Pofut1* (henceforth referred to as Pof cKO), and deletion of both *Eogt* and *Pofut1* (henceforth referred to as EPof dKO), in hematopoietic stem cells. The absolute number of BM cells was significantly increased in both Pof cKO and EPof dKO mice (Figure 5). Gating strategies used to define HSC, HSPC, myeloid and lymphoid cell subsets in BM are shown in supplemental Figure 6. Short-term (ST)-HSC, and LSK (Lineage-Sca1+cKit+) populations were increased in both Pof cKO and EPof dKO mutants, but HSPCs were increased only in EPof dKO BM (Figure 5). By contrast, MPP subsets were reduced except for a small increase in MPP3 cells in EPof dKO BM (Figure 5). Lymphoid primed multipotent progenitor cells (MPP4/LMPP) and common myeloid precursors (CMP) were reduced in Pof cKO and further reduced in EPof dKO BM, and megakaryocyte erythrocyte progenitors (MEP) were decreased in both single and double mutants (Figure 5). By contrast, granulocyte-monocyte progenitors (GMP) were increased in Pof cKO (as observed previously ^22^), and further increased in EPof dKO BM (Figure 5 and supplemental Figure 7). CD11b+/Gr-1+ granulocytes were increased, but common lymphoid progenitors (CLP) were reduced in EPof dKO BM (Figure 5). T-and B-cells in BM were also decreased in EPof dKO mice (Figure 5). NOTCH1 was expressed at the surface of Lin-Sca1+ cells which were increased in frequency in Pof cKO and EPof dKO BM (supplemental Figure 8). Binding of DLL1 and DLL4 Notch ligands was low and similar in control and mutant Sca1+ cells (supplemental Figure 8).

**Figure 4.**
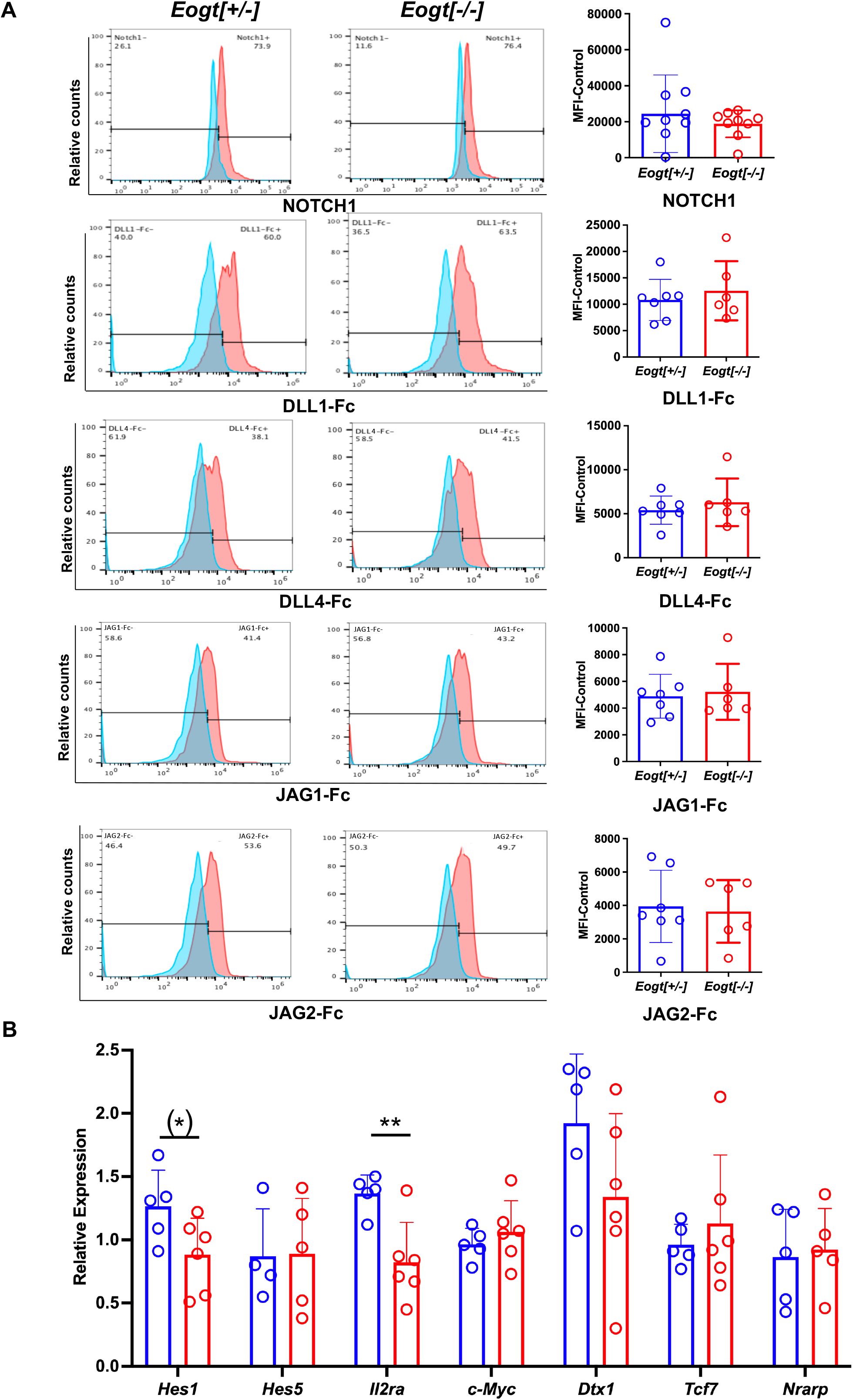
Notch signalling in *Eogt* null T cell progenitors. Representative flow cytometry profiles and histogram quantification of (A) Cell surface NOTCH1, or binding of DLL1-Fc, DLL4-Fc, JAG1-Fc and JAG2-FC to fixed CD4/CD8 DN T cell progenitors from *Eogt*[+/-] or *Eogt*[-/-] mice. Mean fluorescence index (MFI) for anti-Fc Ab was subtracted from MFI for Notch ligand or NOTCH1 Ab (MFI-control). Symbols represent *Eogt*[+/-] (blue circles) and *Eogt*[-/-] (red circles) DN T cells. Fixed cells had been stored for up to 3 months at 4°C. (B) Transcripts from DN T cell progenitors of *Eogt*[+/-] or *Eogt*[-/-] mice were subjected to qRT-PCR as described in Materials and Methods. Relative expression was determined based on the average delta Ct obtained for *Gapdh* and *Hprt* combined. Each symbol represents a mouse of 7-8 weeks. Data are presented as mean ± SEM. *p <0.05, **p<0.01 based on two-tailed Student t test or (*) p <0.05 based on one-tailed Student t test.

**Figure 5.**
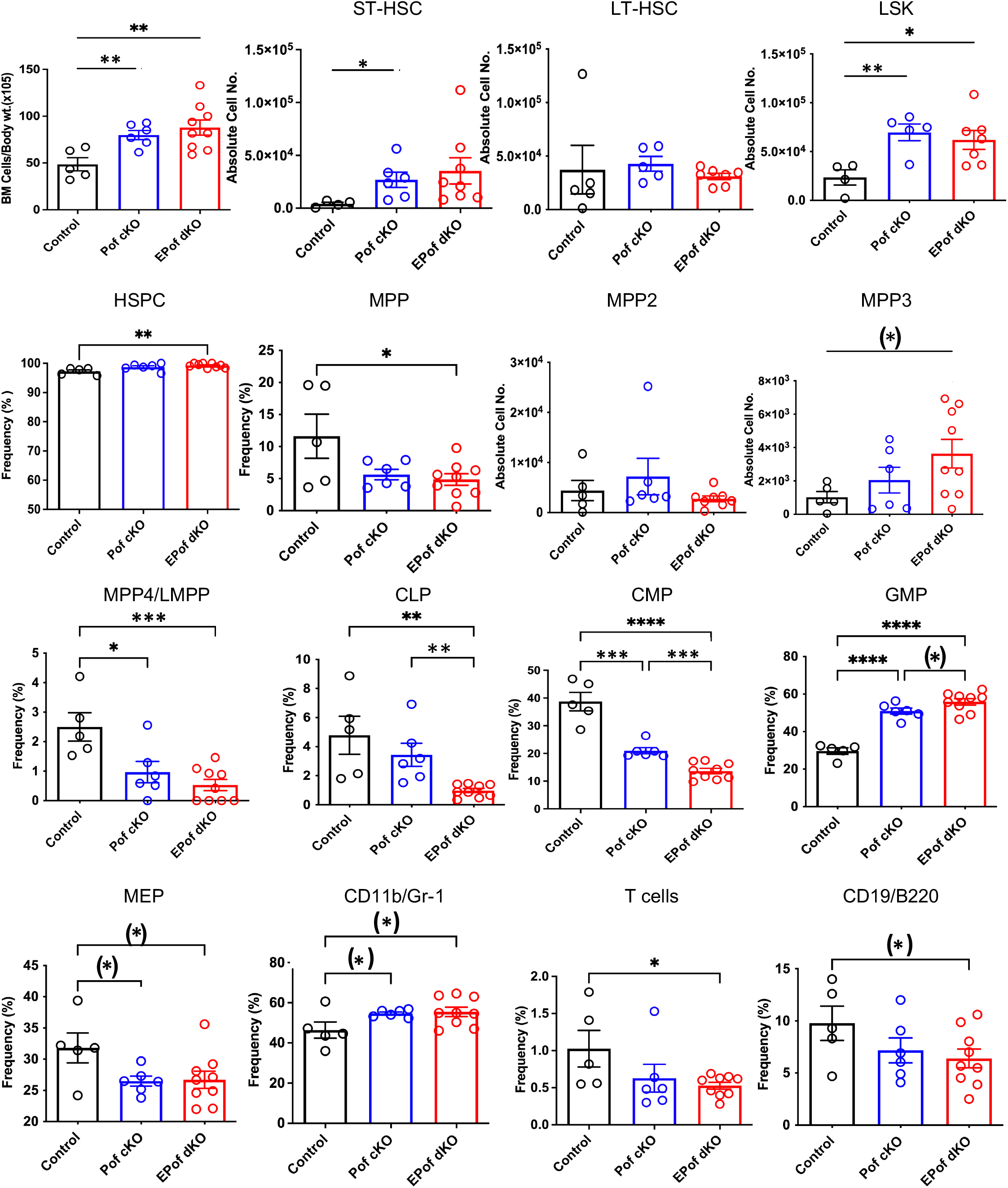
Hematopoiesis in EPof dKO bone marrow. Hematopoiesis was analyzed in bone marrow of *Eogt*[+/-]*Pofut1*[F/F] (Control), *Pofut1*[F/F]:*Vav1-iCre* (Pof cKO) and *Eogt*[-/-]*Pofut1*[F/F*]:Vav1-iCre* (EPof dKO) mice. Absolute numbers or frequency (%) of hemopoietic cell subsets in bone marrow are shown. Gating strategies are shown in supplemental Figure 6 and additional data are shown in supplemental Figure 7. Short term-hemopoietic stem cells (ST-HSC), long-term hemopoietic stem cell (LT-HSC), Lin-Sca1+c-Kit+ (LSK) cells, hematopoietic stem progenitor cells (HSPCs), multipotent progenitor-2, -3, -4 (MPP2, -3, -4), lymphoid primed multipotent progenitor cell (LMPP), common lymphoid progenitor (CLP), common myeloid progenitor (CMP), granulocyte monocyte progenitor (GMP), megakaryocyte erythrocyte progenitor (MEP), T cells, B cells and myeloid cells. Each symbol represents a mouse of 7-8 weeks. Data are presented as mean ± SEM. *p <0.05, **p<0.01, ***p<0.001, ****p<0.001 based on two-tailed Student t test or (*) p <0.05 based on one-tailed Student t test.

### Highly disrupted development of T cells in EPof dKO thymus

Deletion of *Pofut1* in HSC via *Vav1-iCre* led to a marked decrease in thymus weight and a similar reduction was observed in EPof dKO thymus (Figure 6A). The reduced size was accompanied by a dramatic change in T cell maturation (Figure 6B). Early thymic progenitors (ETP) were greatly decreased, and each DN T cell progenitor population (DN1 to DN4) was reduced in absolute cell number in Pof cKO, and even further reduced in EPof dKO thymocytes (Figure 6C and supplemental Figure 9). Interestingly, however, the proportion of DN4 T cells in both single and double mutant thymocytes was increased relative to control (supplemental Figure 9), as observed in thymus lacking all Fringe activities ^36^. The DN thymic population, which includes few if any DN T cell progenitors and all non-T cell populations, was greatly increased in Pof cKO and EPof dKO. This reflected loss of Notch signaling leading to the generation of B cell, myeloid and NK cell subsets (Figure 6D and supplemental Figure 9). However, effects were more severe in EPof dKO than Pof cKO thymus, suggesting that the loss of Notch signaling was greater in EPof dKO thymus.

**Figure 6.**
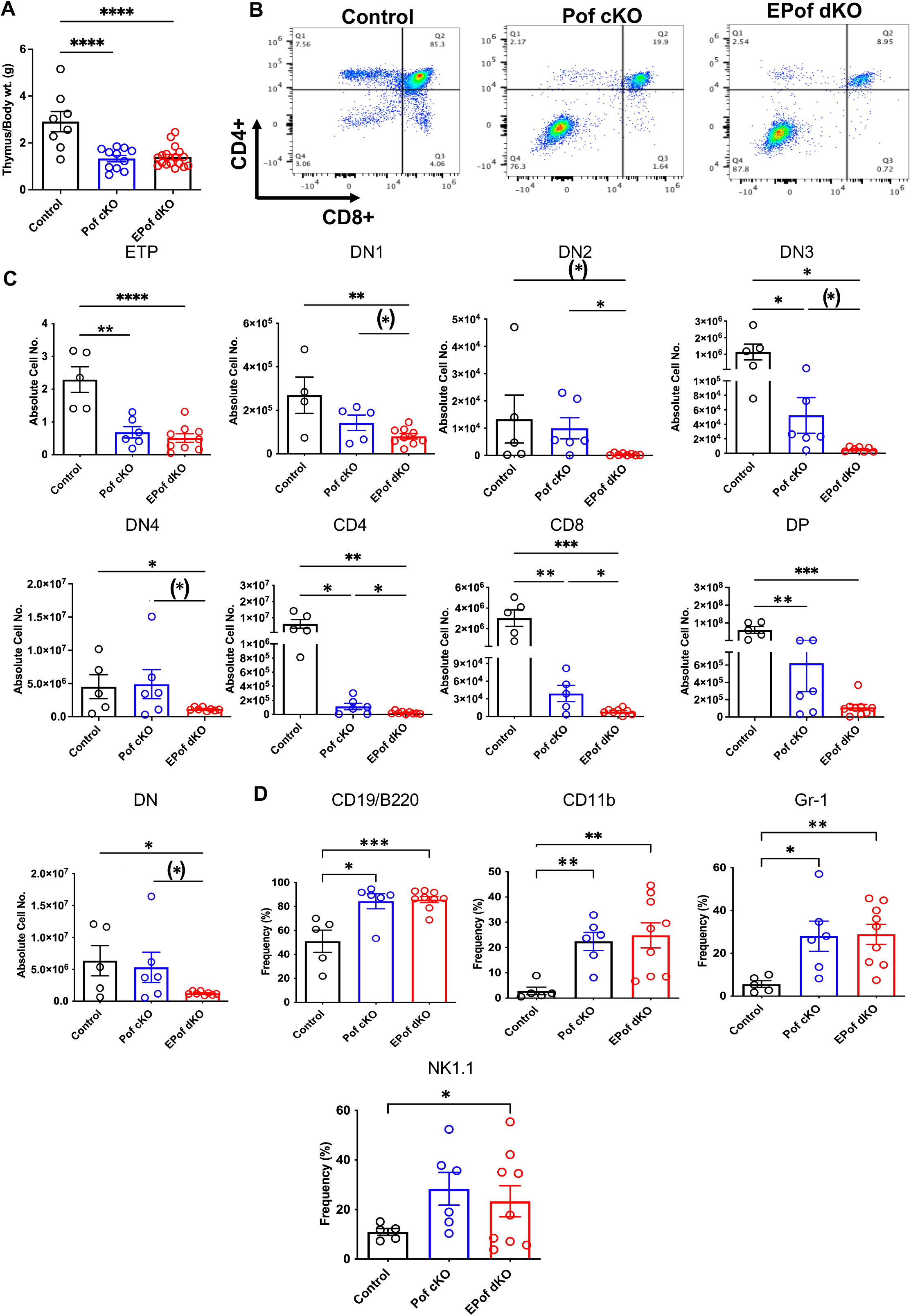
Impaired T cell development in Pof cKO thymus is worse in EPof dKO thymus. (A) Thymus weight compared to body weight. (B) Representative flow cytometry profiles of thymocytes using Abs to CD4 and CD8 cell surface markers after gating on live cells. (C) Absolute cell numbers or frequency (%) of T cell subsets that differed between Control and either mutant. ETP (DN1 cells that were cKit/CD117+), DN1 (CD44+CD25-), DN2 (CD44+CD25+), DN3 (CD44-CD25+), and DN4 (CD44-CD25-) T cell progenitors, DN (CD4 and CD8 double negative), DP (CD4 and CD8 double positive). (D) Absolute cell numbers or frequency (%) of B cell and myeloid cell subsets that differed from control are shown. Additional data are shown in supplemental Figure 9. Each symbol represents a mouse of 7-8 weeks. Data are presented as mean ± SEM. *p <0.05, **p<0.01, ***p<0.001, ****p<0.001 based on two-tailed Student t test or (*) p <0.05 based on one-tailed Student t test.

### Defective B, T and myeloid cell development in EPof dKO spleen

Both Pof cKO and EPof dKO mice had an enlarged spleen (Figure 7A). Absolute numbers of splenocytes and spleen weight were increased in Pof cKO, and further increased in EPof dKO mice (Figure 7B). Histological analysis revealed extramedullary hematopoiesis in both Pof cKO and EPof dKO spleen, with larger areas of extramedullary hematopoiesis observed in EPof dKO spleen (Figure 7A).

The proportion of CD4+ T and CD8+ T cells was significantly reduced in Pof cKO, and further reduced in EPof dKO spleen (Figure 7C and supplemental Figure 10). While CD19+ and B220+ B cells were increased in both Pof cKO and EPof dKO spleen, the CD19+/B220+ B cell population was reduced (Figure 7D). The frequency of Fo-B and the number of MZ-B cells were also reduced (Figure 7D and supplemental Figure 10). While the absolute number of MZ-P cells did not change in the mutants, the relative frequency of MZ-P precursors increased in EPof dKO spleen (Figure 7 and supplemental Figure 10). Myeloid cell subsets such as dendritic cells (CD11b/c+) and Gr-1+ granulocytes were increased in a greater proportion in EPof dKO compared to Pof cKO spleen. Natural killer T cells were reduced in frequency in both Pof cKO and EPof dKO spleen (Figure 7D and supplemental Figure 10).

**Figure 7.**
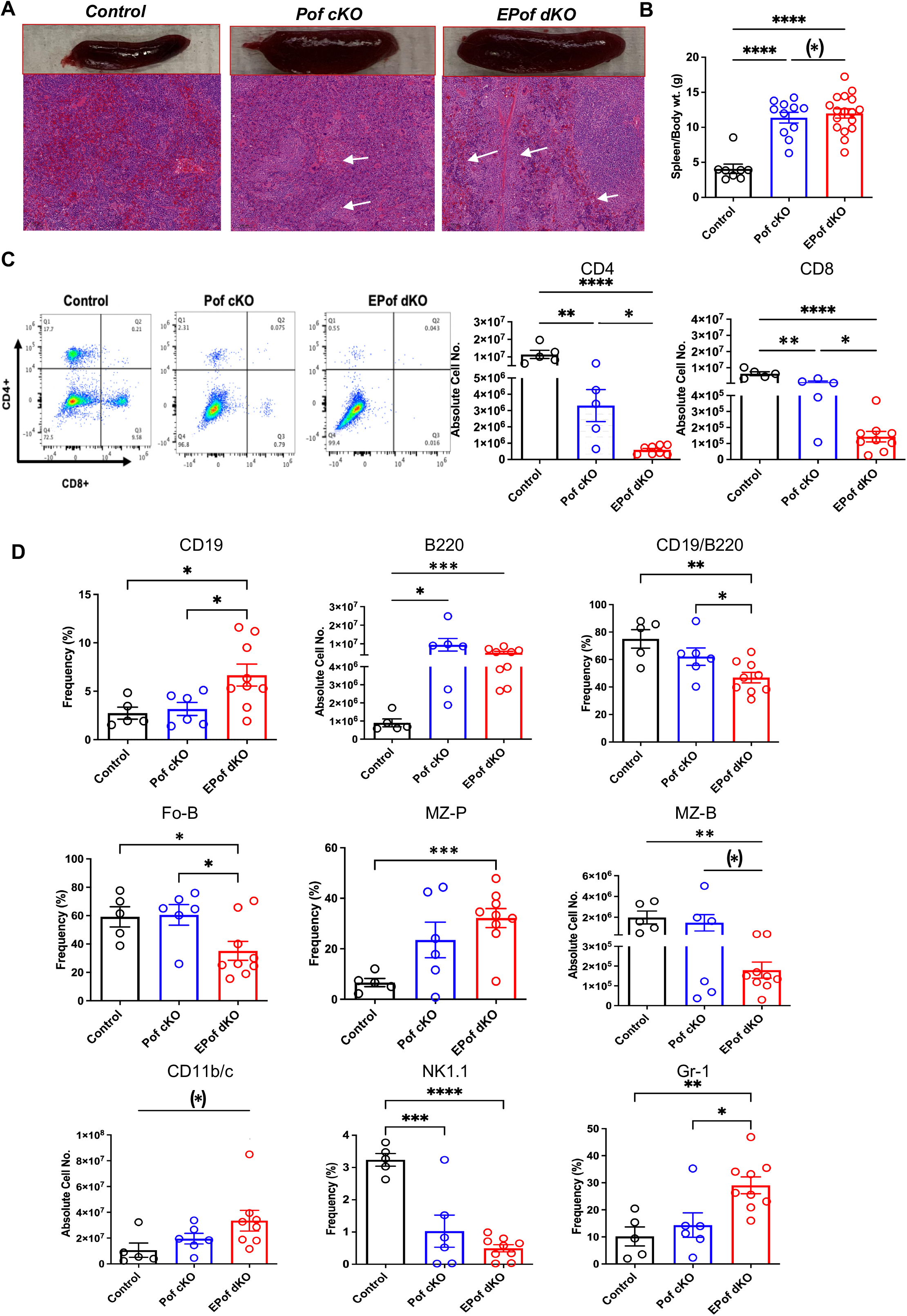
Impaired B cell development in Pof cKO spleen is worse in EPof dKO spleen. (A) Splenomegaly and extramedullary hematopoiesis (original magnification ×10; Bars represent 100 μm in H & E images). (B) Spleen weight compared to body weight. (C) Flow cytometric profiles and histogram quantification for CD4 versus CD8 cell surface expression after gating on live cells. (D) Absolute cell numbers or frequency (%) of splenic lymphoid and myeloid cell subsets that differed from control are shown. CD19/B220 (B cells), Fo-B (follicular B cells), MZ-B (marginal zone-B), MZ-P (marginal zone precursor), CD11b/c (dendritic cells), NK1.1 (natural killer T cells) and Gr-1 (granulocytes). Additional data are shown in supplemental Figure 10. Each symbol represents a mouse of 7-8 weeks. Data are presented as mean ± SEM. *p <0.05, **p<0.01, ***p<0.001, ****p<0.001 based on two-tailed Student t test or (*) p <0.05, based on one-tailed Student t test.

## Discussion

Defining specific roles for the glycans that regulate Notch signaling in lymphoid and myeloid development facilitates our understanding of cell fate decisions regulated by Notch signaling, and of potential consequences for people with congenital diseases that perturb Notch signaling ^2,39^. Several congenital diseases inhibit the synthesis or extension of O-glycans that regulate Notch signaling and may induce immune cell defects. Mutations in *EOGT* cause Adams Oliver syndrome ^40^ and autosomal dominant mutations in *POFUT1* cause Dowling Degos Disease 2 (DDD2) ^41^. Here we show that the generation of certain lymphoid and myeloid subsets in bone marrow, thymus and spleen was perturbed in mice lacking *Eogt*. In bone marrow, loss of EOGT caused increased numbers of B cells and granulocytes. In thymus, the phenotype of *Eogt* null mice was similar, but not identical to, mice which lack all three Fringe genes (*Fng* tKO) ^36^. Early T cell progenitors DN1 and DN2 were reduced in frequency, whereas DN4 progenitors were increased, DP T cells were slightly reduced, and CD4+ and CD8+ SP T cells were increased. Inhibition of Notch signaling in thymus is well known to lead to the generation of B cells and myeloid cells in place of T cells ^9,11,42,43^. *Eogt* null thymus contained significantly increased numbers of B cells and granulocytes, and an increased frequency of myeloid cells. In spleen, *Eogt* null splenocytes included increased numbers of several B cell subsets, although there were no effects on T cells, unlike in *Fng* tKO mice which had a reduced frequency of T cells in spleen ^36^. Deletion of RBP-Jk by *Vav1*-*iCre* results in an increased frequency of CD19+ B cells ^44^, also observed in *Eogt* null spleen. *Eogt* null mice showed increased absolute numbers of CD19+ B cells, increased Fo-B cells, and a decrease in natural killer T-cells and dendritic cells in the spleen. The overall *Eogt* null phenotype was largely cell autonomous following bone marrow transplantation. In addition, expression of Notch target genes *Hes1* and *Il2ra* was reduced, similar to *Fng* tKO DN T cell progenitors that had reduced *Il2ra* and *Dtx1* expression ^36^. The combined data provide strong evidence that EOGT and O-GlcNAc glycans are required for optimal Notch signaling in the development of lymphoid and myeloid cells from HSC.

Further evidence that EOGT and O-GlcNAc glycans support Notch signaling in lymphoid and myeloid development was obtained in compound mutant mice lacking *Eogt* and conditionally lacking *Pofut1* in HSC. EPof dKO lymphoid and myeloid populations in BM, thymus and spleen were more affected compared to Pof cKO. We conclude that the loss of *Eogt* and O-GlcNAc glycans in Pof cKO HSC exacerbated the deficits in T, B and myeloid differentiation evident in Pof cKO mice. This result, and our findings that *Eogt* is required for optimal Notch signaling and the differentiation of HSC, provide an explanation for the observation that *Pofut1*:*Mx1-Cre* T cell deficiencies were not as severe as those obtained in *RBP-Jk:Mx1-Cre* thymus ^23^. In the absence of POFUT1 and O-fucose glycans, the O-GlcNAc glycans transferred by EOGT to Notch receptors can support a low but significant level of Notch signaling. Thus, the O-fucose and O-GlcNAc glycans on Notch act synergistically to provide optimal Notch signaling in lymphopoiesis and myelopoiesis.

The conclusions obtained from these experiments are necessarily limited by the difficulty of demonstrating structural changes in the O-glycans on Notch receptors predicted to change in the absence of relevant glycosyltransferase(s). Thus, to obtain sufficient quantities of NOTCH1 from splenic T cells for analysis by mass spectrometry, it was necessary to activate the T cells which increases cell surface NOTCH1 by ∼ 10-fold ^45^. In addition, the effects of conditional deletion of *Eogt* in HSC would allow us to define the contribution, if any, of *Eogt* null stroma to the *Eogt* null phenotype. It would also be important to determine Notch ligand binding and Notch target gene expression in cells from different HSPC lineages in Pof cKO and EPof dKO mice. Single cell RNA-seq of different mutant HSC and HSPCs would greatly expand our understanding of the pathways affected by altered Notch signaling due to loss of regulation by O-glycans in HSC. Finally, it would be important to investigate roles for the O-glycans on Notch ligands by deleting *Pofut1, Eogt* and related glycosyltransferase genes in stromal cells.

## Acknowledgments

We thank Dr. Jinghang Zhang and Aodengtuya of the Flow Cytometry Core Facility of the Albert Einstein Cancer Center for their assistance in discussing multi-color flow cytometry experimental strategies. We are also thankful to Subha Sundaram for her technical support and Swathi-Rao Narayanagari for her assistance in the bone marrow transplantation (BMT) study. We thank Hillary Guzik and Andrea Briceno from the Analytical Imaging Facility of the Einstein Cancer Center for slide scanning using a 3D Histech P250 High-Capacity Slide Scanner purchased with NIH SIG #1S10OD019961-01.

This work was supported by research funding from the National Institutes of Health grant RO1 GM-106417 (P.S.) and National Institutes of Health grant PO1 grant CA-13330 (partial support from core facilities).

## Authorship

Conceptualization, PS; Methodology, PS, AT; Investigation, AT, PS; Visualization AT, PS; Funding acquisition, PS; Project administration, PS; Supervision, PS; Writing – original draft, AT; review and editing, PS, AT.

### Conflict-of-interest disclosure

Authors declare that they have no competing interests.

## Notes

### Competing Interest Statement

The authors have declared no competing interest.

### Summary of Updates

The manuscript has been revised to include critiques from colleagues and to include new data on Lin-Sca1+ bone marrow cells.

