## Supplemental material for "Synergistic regulation of Notch signaling by different O-glycans promotes hematopoiesis"

**Short Title:** O-glycans promote lymphopoiesis and myelopoiesis

**Corresponding author:**

Pamela Stanley

Department of Cell Biology,

Albert Einstein College of Medicine

New York, NY, 10461 USA

### Supplemental Appendix


| Supplemental Figures | Page |
|---|---|
| **Supplemental Figure 1.** Gating strategy for BM hematopoietic cell subsets. Related to Figure 2, Figure 5 and Supplemental Figure 4 | 3 |
| **Supplemental Figure 2.** Gating strategy for thymic immune cell subsets. Related to Figure 2, Figure 6, Supplemental Figures 4 and 5 | 4 |
| **Supplemental Figure 3.** Gating strategy for splenic immune cell subsets. Related to Figure 2, Figure 7, Supplemental Figures 4 and 5 | 5 |
| **Supplemental Figure 4.** Immune cell development in *Eogt*[+/+] versus *Eogt*[+/-] mice. Related to Figure 2 | 6 |
| **Supplemental Figure 5.** Immune cell subsets from Control versus *Eogt*[-/-] mice. Related to Figure 2 | 7 |
| **Supplemental Figure 6.** Gating strategy for BM hematopoietic cell subsets. Related to Figure 5 | 8 |
| **Supplemental Figure 7.** Hematopoietic stem progenitors in control, Pof cKO and EPof dKO BM. Related to Figure 5 | 9 |
| **Supplemental Figure 8.** Notch ligand binding to Lin-Sca1+ cells. Related to Figure 5 | 10 |
| **Supplemental Figure 9.** Thymic immune cell subsets in Pof cKO and EPof dKO mice. Related to Figure 6 | 11 |
| **Supplemental Figure 10.** Splenic immune cell subsets in Pof cKO and EPof dKO mice. Related to Figure 7 | 12 |

| Supplemental Tables | |
|---|---|
| **Supplemental Table 1.** Primer sequences used in genotyping and qRT-PCR | 13 |
| **Supplemental Table 2.** Antibodies used in this work | 14-15 |

### Bone Marrow

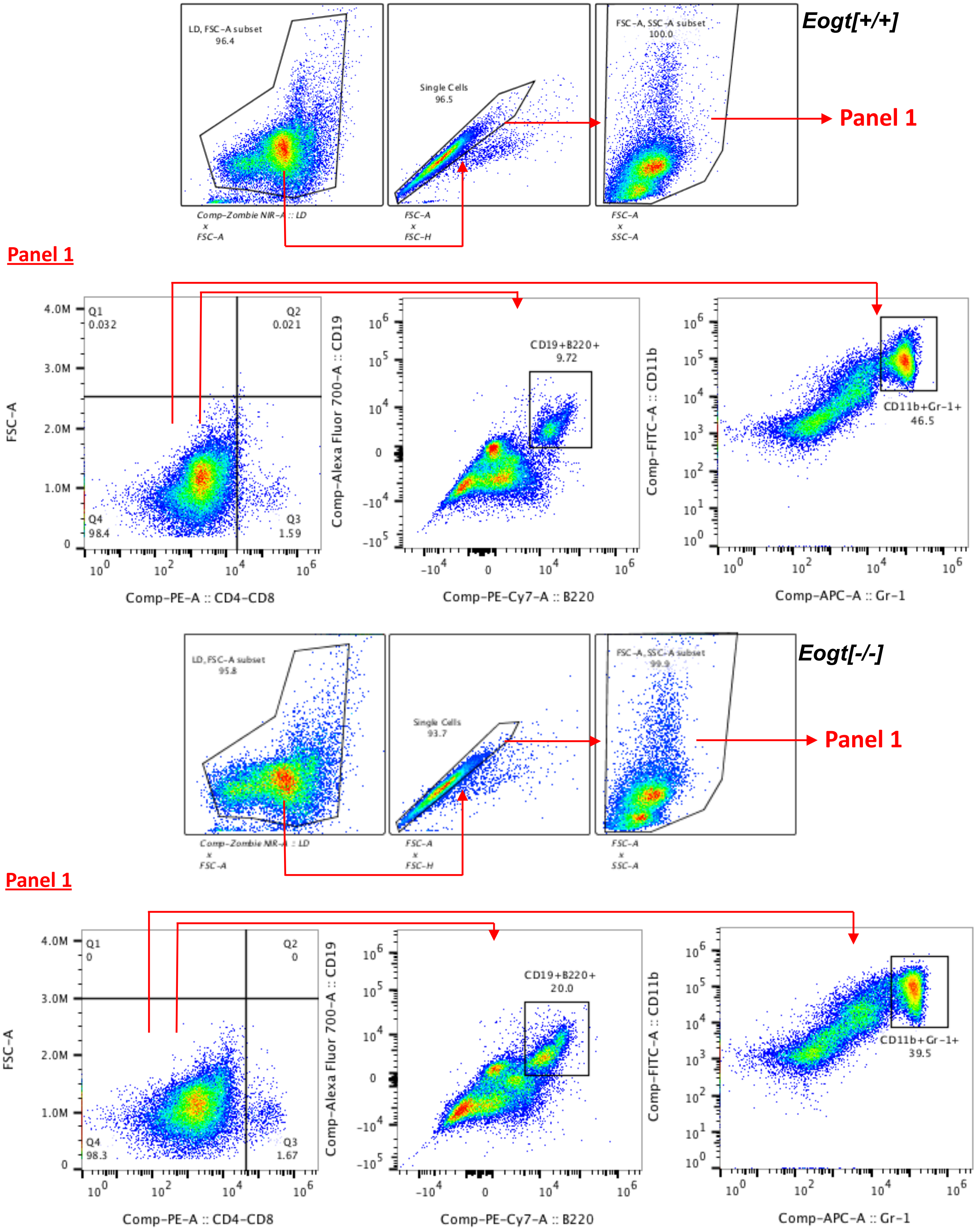


**Supplemental Figure 1. Gating strategy for BM immune cell subsets.** Related to Figure 2, Figure 5, and Supplemental Figure 4

Flow cytometry profiles of BM cells from a 7-8 week *Eogt*[+/+] or *Eogt*[-/-] mouse selected after gating on live cells using Zombie NIR dye in the Cytek$^{TM}$ Aurora flow cytometer. Live singlets were selected (FSC-H vs FSC-A) and the main population (SSC-A vs FSC-A) was plotted for CD4+CD8+ vs FSC-A, and gated on CD4-CD8- cells to compare CD19+ vs B220+ or CD11b+ vs Gr-1+ cell populations.

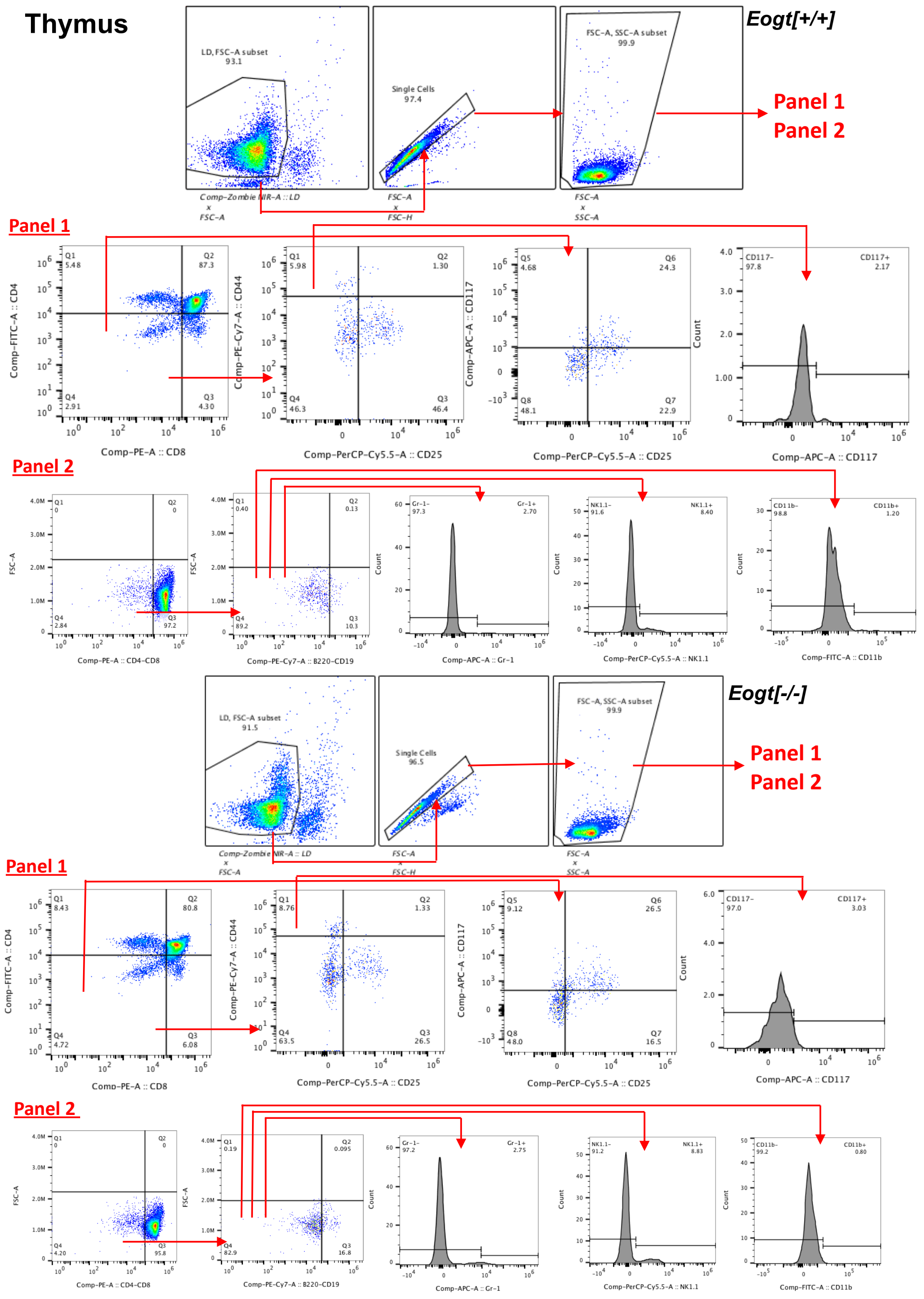


**Supplemental Figure 2. Gating strategy for thymic immune cell subsets.** Related to Figure 2, Figure 6 and Supplemental Figures 4 and 5

Flow cytometry profiles of thymocytes from a 7-8 week *Eogt*[+/+] or *Eogt*[-/-] mouse selected after gating on live cells using Zombie NIR dye in the Cytek$^{TM}$ Aurora flow cytometer. Live singlets were selected (FSC-H vs FSC-A) and the main population (SSC-A vs FSC-A) was plotted for Panel 1 antibodies to CD4 vs CD8 to determine frequencies and absolute numbers of CD4+ SP and CD8+ SP T cells, CD4+CD8+ DP T cells and CD4-CD8- DN T cells. The DN T cell population was plotted for CD44 vs CD25 to determine frequencies and absolute numbers of CD44+CD25- (DN1), CD44+CD25+ (DN2), CD44-CD25+ (DN3) and CD44-CD25- (DN4) T cell progenitors, and CD117+ DN1 cells (ETP). Panel 2 antibodies were used to plot FSC-A vs CD4/CD8 to gate on CD4-CD8- cells to then determine frequencies and absolute numbers of non-T cells in the thymus by plotting CD19 vs B220, and gating on CD19-B220- to determine non-B Gr-1+, NK1.1+, CD11b+ cells. Flow cytometry plots show B cells (B220+/CD19+), myeloid cells (CD11b+), granulocytes (Gr-1+), and natural killer T-cells (NK1.1+).

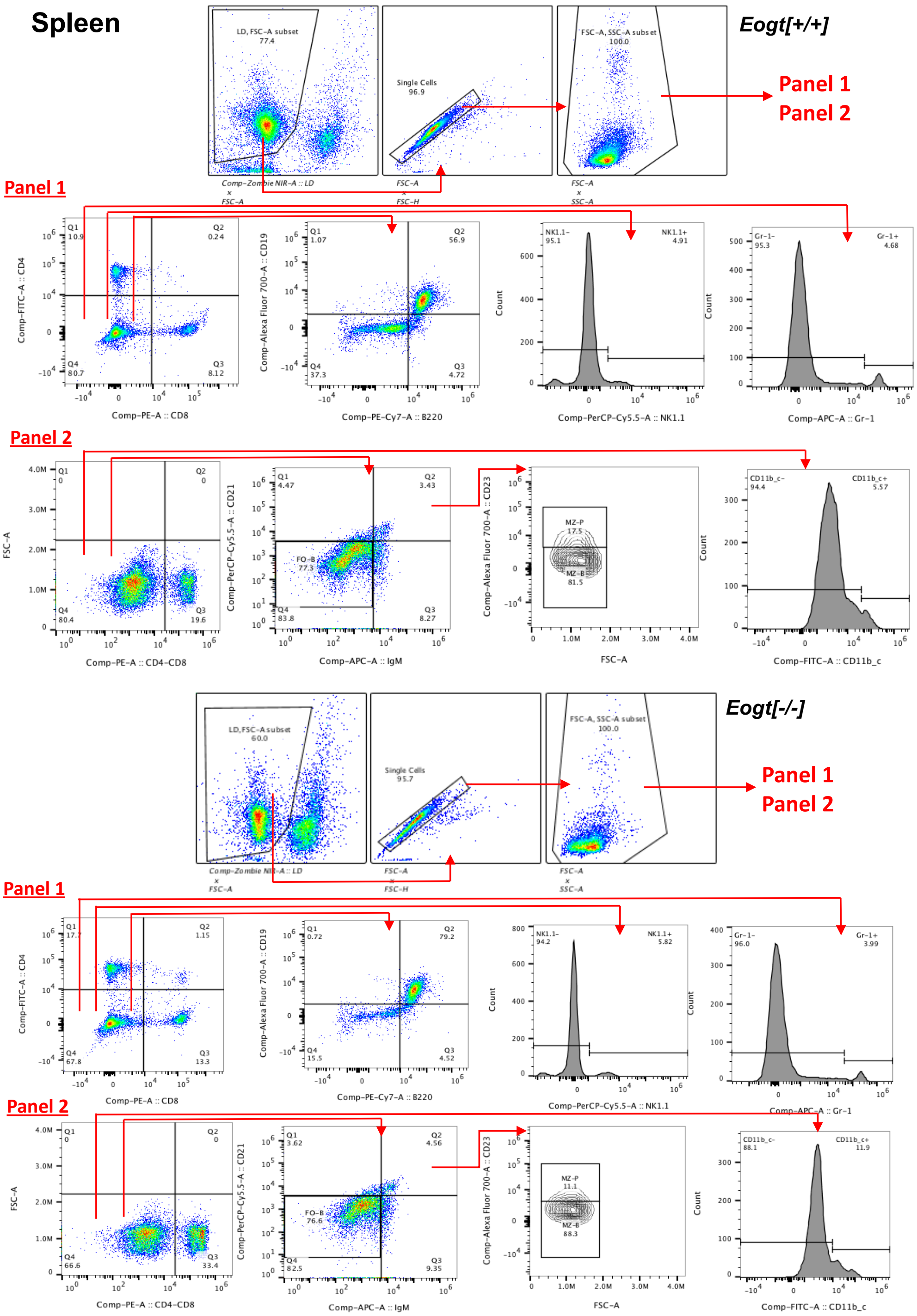


**Supplemental Figure 3. Gating strategy for splenic immune cell subsets.** Related to Figure 2, Figure 7 and Supplemental Figure 4 and 5.

Flow cytometry profiles of splenocytes from a 7-8 week *Eogt*[+/+] or *Eogt*[-/-] mouse selected after gating on live cells using Zombie NIR dye in the Cytek$^{TM}$ Aurora flow cytometer. Live singlets were selected (FSC-H vs FSC-A) and the main population (SSC-A vs FSC-A) was used with Panel 1 antibodies to plot CD4 vs CD8**;** non-T cells (CD4-CD8-) were used to plot CD19 vs B220 to determine frequencies and absolute numbers of B cells, natural killer T cells (NK1.1), or granulocytes (Gr-1)**.** Panel 2 antibodies were used with the main population of singlets to plot FSC-A vs CD4/CD8**;** non-T cells (CD4-/CD8-) were used to plot CD21 vs IgM or CD11b/c**;** CD21+IgM+ were used to plot CD23+ vs FSC-A. Flow cytometric profiles show Fo-B cells (IgM $^{int/lo}$ CD21+) and histogram showing CD11b. The IgM+CD21+ subset was further subdivided into MZ-P and MZ-B cells based on forward scatter and CD23 expression.

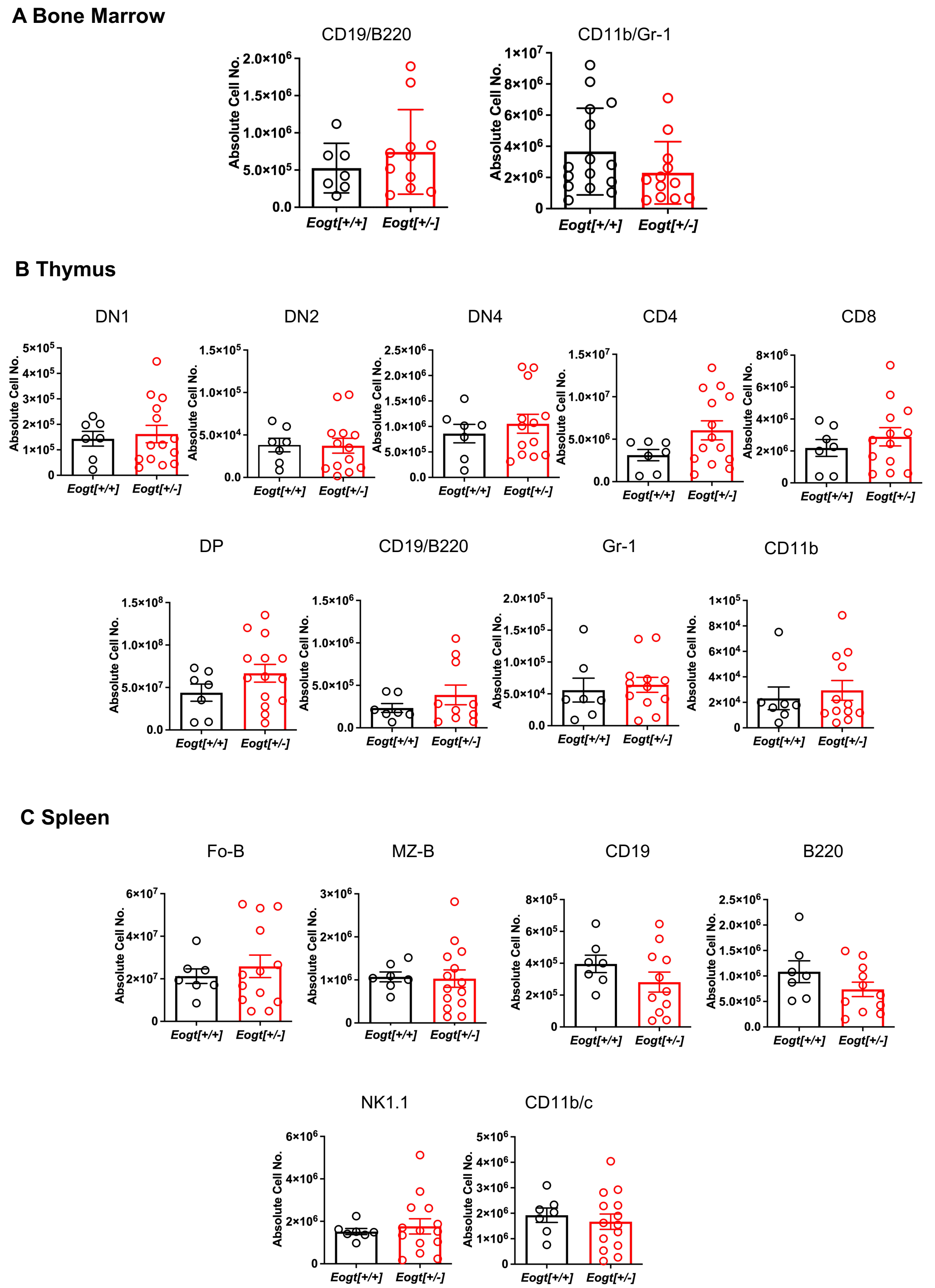


**Supplemental Figure 4. Immune cell development in *Eogt* [+/+] and *Eogt*[+/-] mice.** Related to Figure 2

(A) Absolute cell numbers of CD19+ B220+ B cells, and granulocytes (CD11b+Gr-1+) in bone marrow. (B) Absolute cell numbers of CD44+CD25- (DN1), CD44+CD25+ (DN2), and CD44-CD25- (DN4) T cell progenitors, CD4+ SP, CD8+ SP, CD4+CD8+ DP T cells and B cells (B220+/CD19+), granulocytes (Gr-1+) and myeloid cells (CD11b+) in thymus. (C) Absolute numbers of follicular-B (Fo-B), marginal zone B (MZ-B), CD19+, B220+ B cells, natural killer T cells (NK1.1+), and dendritic cells (CD11b/c+). Each symbol represents a mouse of 7-8 weeks.

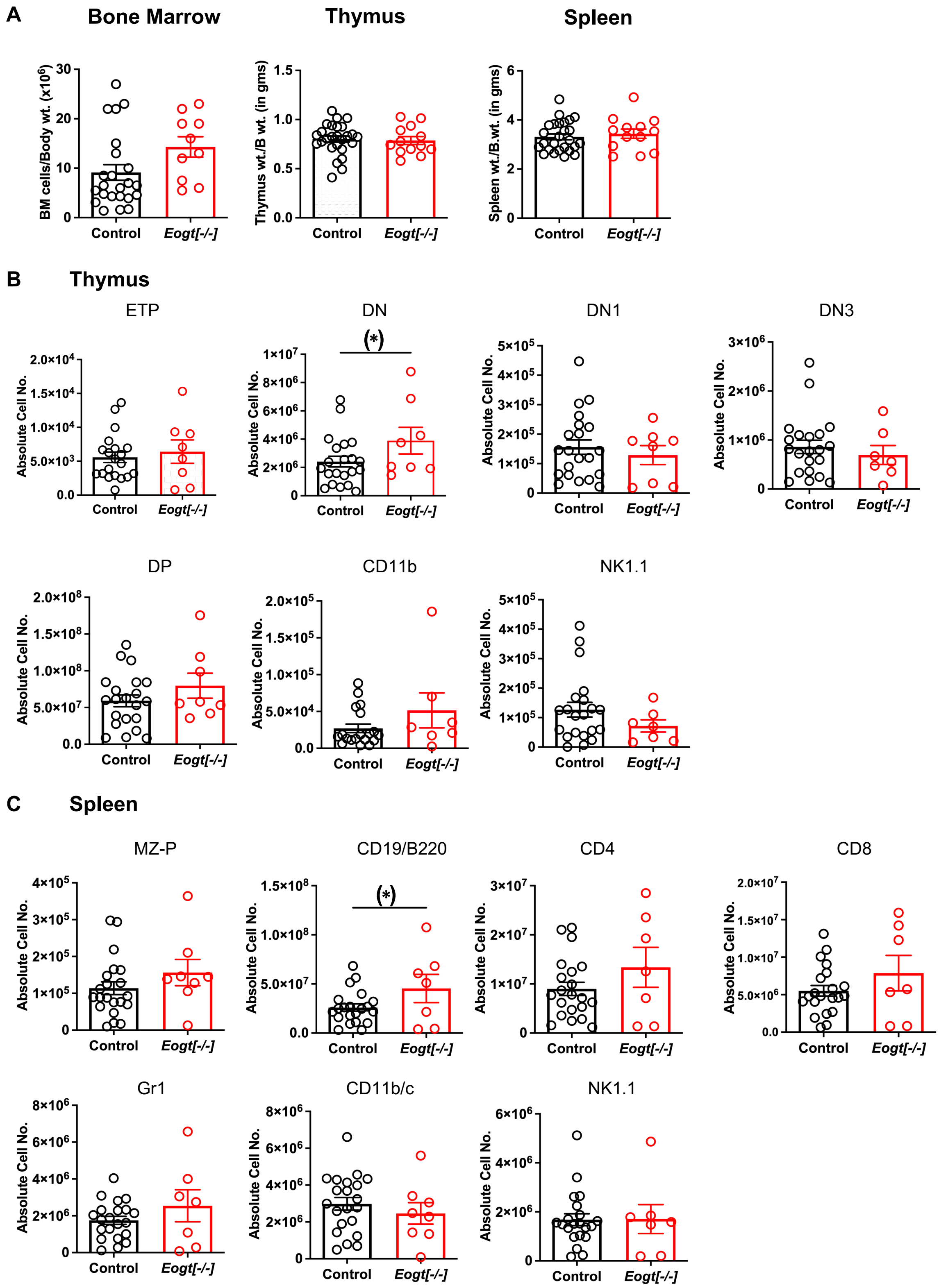


**Supplemental Figure 5. Immune cell subsets in Control versus *Eogt*[-/-] mice.** Related to Figure 2.

(A) Bone Marrow cellularity, thymus to body weight and spleen to body weight. (B) Absolute cell numbers of ETP, DN, DN1, DN3, DP, myeloid cells (CD11b+) and natural killer T cells (NK1.1+) in thymus. (C) Absolute cell numbers MZ-B, B cells (CD19+/B220+), CD4+, CD8+ T cells, granulocytes (Gr-1+), dendritic cells (CD11b/c), and natural killer T cells (NK1.1) in spleen. Each symbol represents a mouse of 7-8 weeks. Data presented as mean SEM. (*) p <0.05 based on one-tailed Student t test.

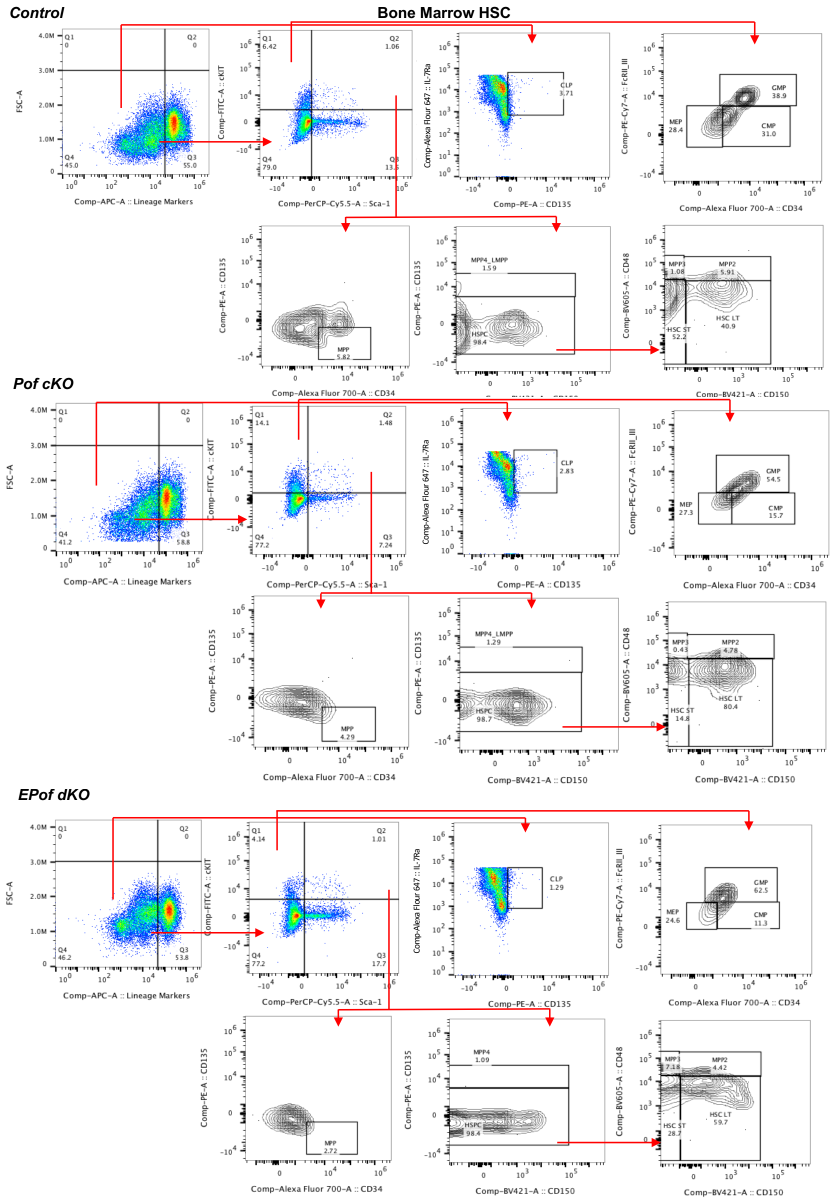


**Supplemental Figure 6. Gating strategy for BM hematopoietic cell subsets.** Related to Figure 5.

Flow cytometric profiles of targeted cell populations were selected after gating on live cells using Zombie NIR dye using the Cytek$^{TM}$ Aurora flow cytometer from 7-8 weeks old, Control, Pof cKO and EPof dKO. HSPC populations were identified as follows:

LSK: Lin- cKit+ Sca1+,

Long-term hematopoietic stem cells (LT-HSC); Lin−Sca1+Kit+ CD150+CD135−,

Short-term hematopoietic stem cells (ST-HSC); Lin−Sca1+Kit+ CD150−CD135−,

Multipotent progenitors (MPP); Lin−Sca1+Kit+ CD150−CD135+,

Lymphoid primed multipotent progenitor cells (LMPP); Lin−Sca1+Kit+ CD135$^{high}$,

Common lymphoid progenitors (CLP); Lin−KitintCD135+IL7Ra+,

Granulocyte/macrophage progenitors (GMP), Lin−Kit+Sca−FcRII/III+CD34+,

Megakaryocyte's erythrocytes progenitors' cells (MEP); Lin−IL-7R−c-kit+Sca-1−CD34−FcRII/III$^{low}$

Common myeloid progenitors (CMP); Lin−Kit+Sca−FcRII/III−CD34+,

Multipotent Progenitors cells 2 (MPP2); Lin−Kit+Sca+CD150+CD48+,

Multipotent Progenitors cells 3 (MPP3); Lin−Kit+Sca+CD150-CD48+

Multipotent Progenitors cells (MPP4); Lin−Kit+Sca+CD135+CD150-

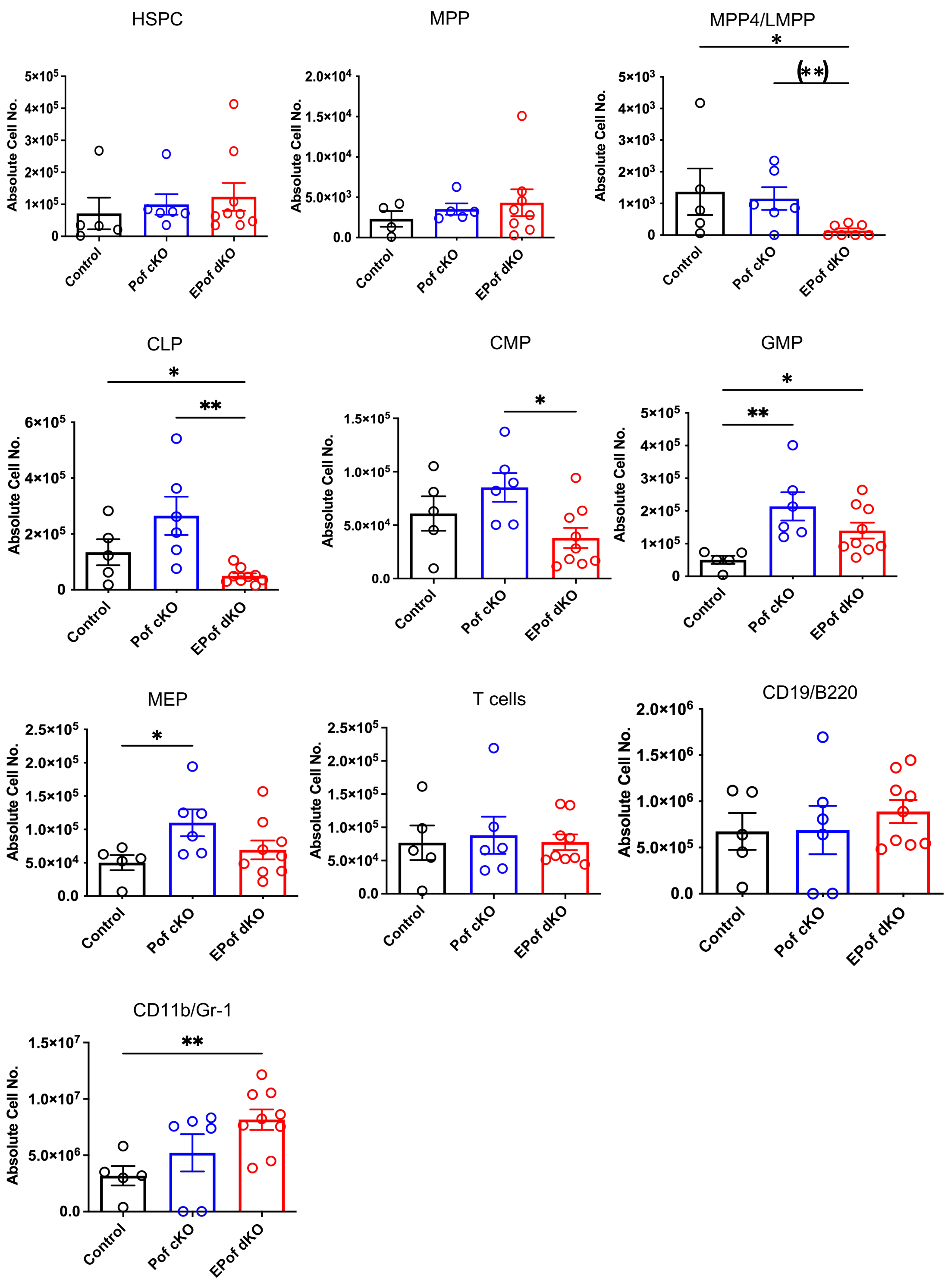


**Supplemental Figure 7. Hematopoietic stem progenitors in Pof cKO and EPof dKO BM.** Related to Figure 5.

Hematopoiesis, lymphoid and myeloid cell development were compared in Pof cKO versus EPof dKO versus Control mice with no *Vav1*-iCre. Each symbol represents a mouse of 7-8 weeks. Absolute cell numbers of different hematopoietic stem progenitor cells (HSPCs); multipotent progenitor (MPP) and multipotent progenitor-4 (MPP4)/LMPP cells, common lymphoid progenitor (CLP) common myeloid progenitor (CMP), granulocytes monocytes progenitor (GMP), megakaryocytes erythrocytes progenitors (MEP), T cells and B cells (B220/CD19) and granulocytes (CD11b/Gr-1) subsets.

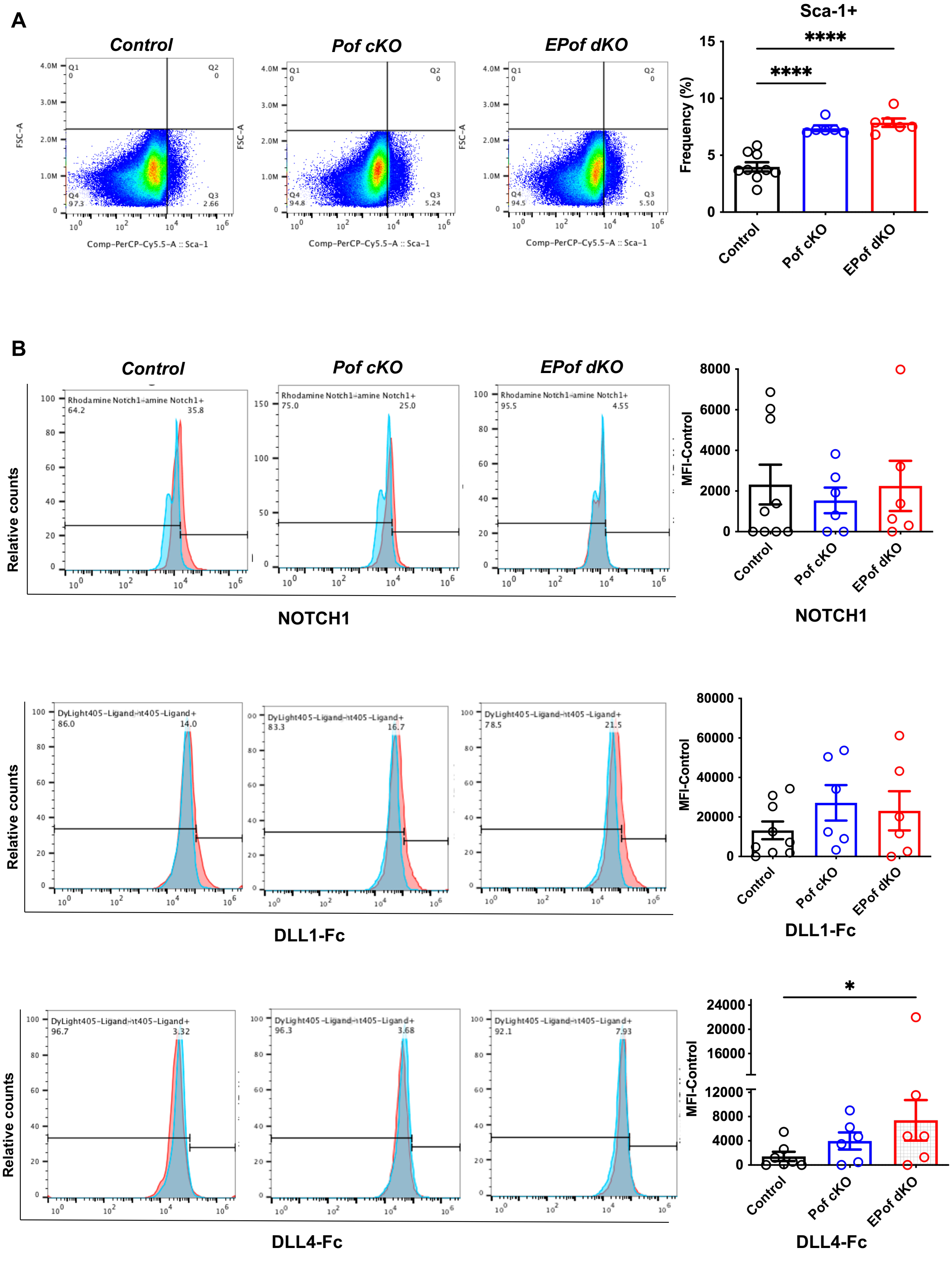


**Supplemental Figure 8. Notch ligand binding to Lin-Sca1+ cells.** Related to Figure 5.

Gating of Lin-Sca1+ cells and frequency (%) in triplicate samples of BM from Control (n=3), Pof cKO (n=2) and EPof dKO (n=2) 6-7 weeks old mice. All replicates are shown. Flow cytometry profiles of NOTCH1 cell surface expression, and binding of DLL1-Fc or DLL4-Fc to Lin-Sca1+ cells from the same mice performed in triplicate.

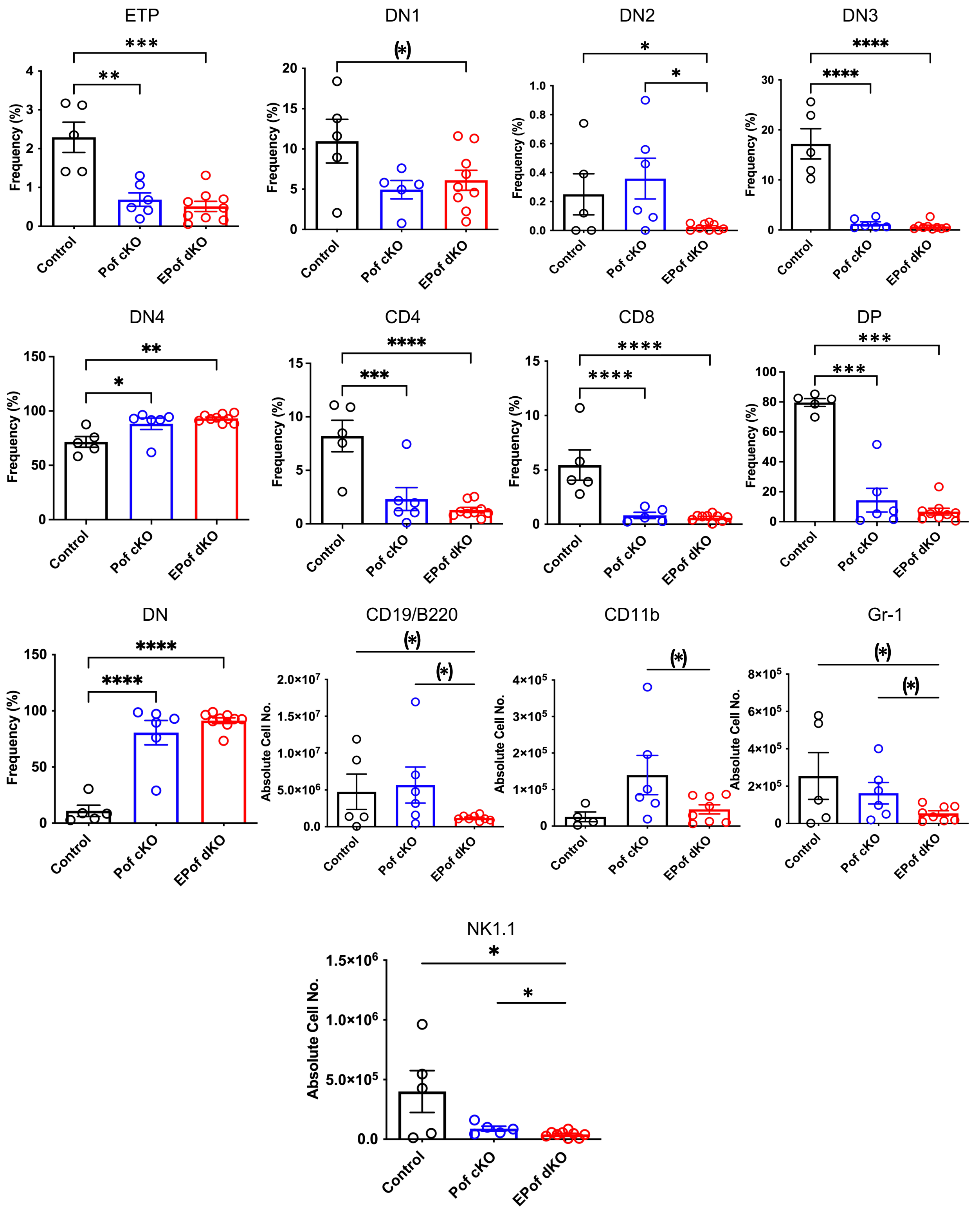


**Supplemental Figure 9. Thymic immune cell subsets in Pof cKO and EPof dKO mice.** Related to Figure 6.

Frequency(%) of different T-cell subsets; ETP(DN1 cells that were cKit/CD117+), and stages of different double negative (DN) T cells; CD44+CD25- (DN1), CD44+CD25+ (DN2), CD44-CD25+ (DN3), and CD44-CD25- (DN4) T cell progenitors, CD4, CD8, Double Positive (DP), Double negative (DN), and absolute numbers of B cells (B220/CD19), myeloid cells (CD11b), granulocytes (Gr-1) natural killer T-cells (NK1.1). Each symbol represents a mouse 7-8 weeks. Data are presented as mean SEM. *p <0.05, **p<0.01,***p<0.001, ,****p<0.0001 based on two-tailed Student t test and (*)p <0.05 based on one-tailed Student t test.

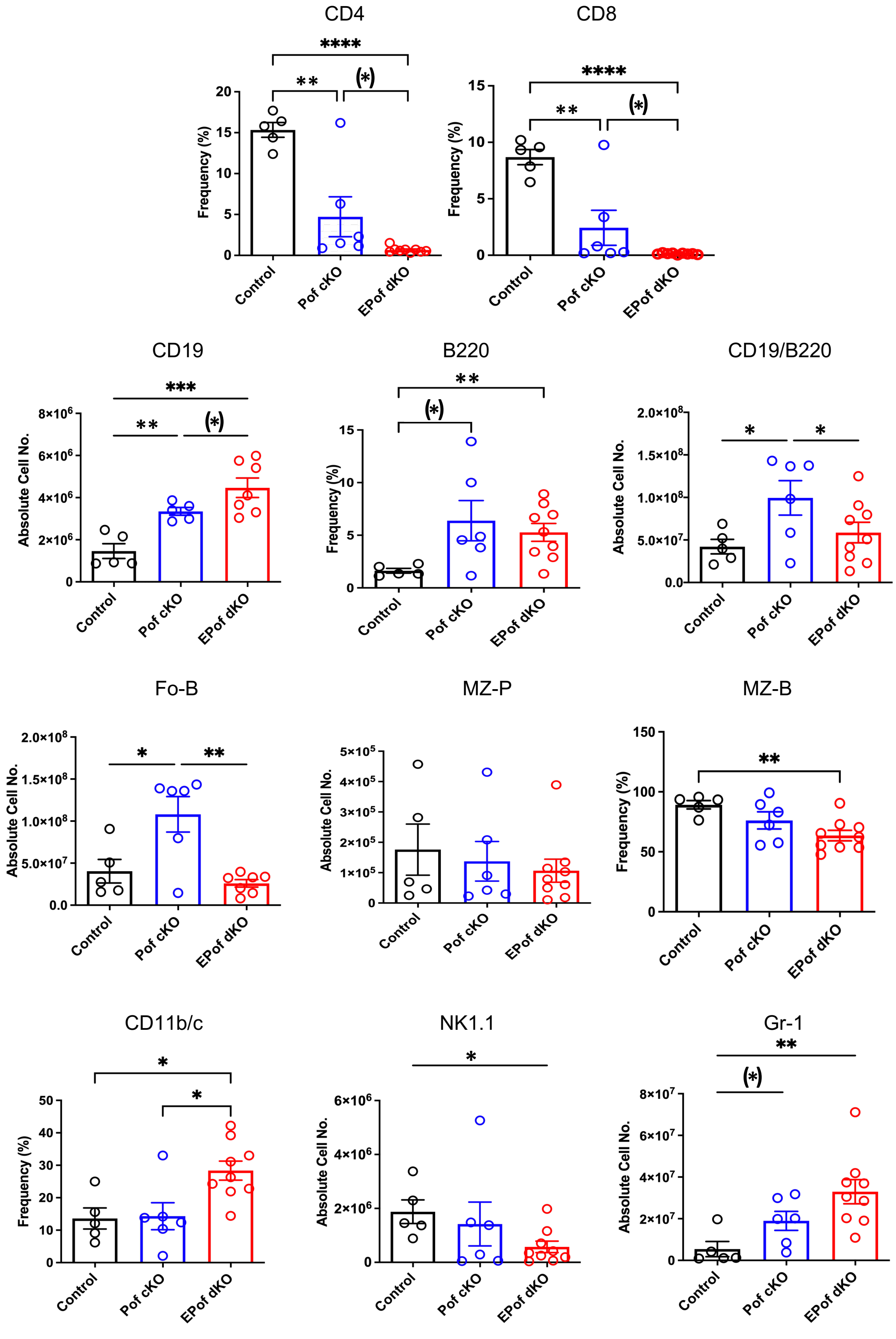


**Supplemental Figure 10. Splenic immune cell subsets in Pof cKO and EPof dKO mice.** Related to Figure 7.

Absolute cell numbers of CD19, B-cells (B220/CD19), follicular B cells (Fo-B), marginal zone precursors (MZ-P) cells, natural killer T cell (NK1.1) and granulocytes (Gr-1). Frequency (%) of splenic T-cell subsets; CD4, CD8, B220, marginal zone-B (MZ-B), dendritic cells (CD11b/c). Each symbol represents a mouse 7-8 weeks. Data are presented as mean SEM. *p <0.05, **p<0.01,***p<0.001, ,****p<0.0001 based on two-tailed Student t test and (*)p <0.05 based on one-tailed Student t test.

**Supplemental Table 1. Primer sequences used in genotyping and qRT-PCR**

| Gene | Name | Sequence |
|---|---|---|
| *Pofut1 F/del* | PS644/Forward | *GGGTCACCTTCATGTACAAGTGAGTG* |
| | PS645/Reverse | *ACCCACAGGCTGTGCAGTCTTTG* |
| *Pofut1 F/wt* | FB21/Forward | *CCAGGCTGATCACTTCTTGG* |
| | FB22/Reverse | *CCCTGTCTCGAAAAAGCAAA* |
| *Eogt* | 3rd loxF/Forward | *CCACCCGACCCCTGCCAGAACATAATGCTCTCTTGCATC* |
| | 3rd loxR/Reverse | *GCTGTCGCCAGAGGAGAGAGTGGGTGCTTACTTAC* |
| | 25307 Rv/Reverse | *CCAAGGCGGTCTTGGCCCAT* |
| *Vav1-icre* | FW internal/Forward | *CTAGGCCACAGAATTGAAAGATCT* |
| | RV Internal/Reverse | *GTAGGTGGAAATTCTAGCATCATCC* |
| | FW Transgene/Forward | *AGATGCCAGGACATCAGGAACCTG* |
| | RV Transgene/Reverse | *ATCAGCCACACCAGACACAGAGATC* |
| *Hes1* | Forward | *AAGGCAGACATTCTGGAAAT* |
| | Reverse | *GTCACCTCGTTCATGCACTC* |
| *Hes5* | Forward | *GGACCAGAGGATGAGCTCGTT* |
| | Reverse | *AGGAGGGAGCCTTCGGAAGA* |
| *cMyc* | Forward | *AGTGCTGCATGAGGAGACAC* |
| | Reverse | *GGTTTGCCTCTTCTCCACAG* |
| *Deltex1* | Forward | *CATCAGTTCCGGCAAGAC* |
| | Reverse | *ATGGTGATGCAGATGTCC* |
| *CD25* | Forward | *GGAATTGGTCTATATGCGTTGCTTA* |
| | Reverse | *CATGTCTGTTGTGGTTTGTTGCTCT* |
| *Tcf7* | Forward | *TCCGAGTACATGGAGAAGCC* |
| | Reverse | *GGGTAGGGCATGAGCAGATT* |
| *Nrarp* | Forward | *AGATACCGGATCCTCCGCTT* |
| | Reverse | *CCGGTAATGGTTGTTTGGCG* |
| *Gapdh* | Forward | *AAGGTCATCCCAGAGCTGAA* |
| | Reverse | *CTGCTTCACCACCTTCTTGA* |
| *Hprt* | Forward | *GGACCTCTCGAAGTGTTGGATAC* |
| | Reverse | *GCTCATCTTAGGCTTTGTATTTGGCT* |

**Supplemental Table 2. Antibodies used in this work**

| Antibody | Fluorochrome | Isotype | Clone | Cat# | Company |
|---|---|---|---|---|---|
| **CD4** | PE | Rat IgG2a K | RM4.5 | 553048 | BD Pharmingen |
| **CD8a (ly2)** | PE | Rat IgG2a K | 53-6.7 | 553033 | BD Pharmingen |
| **CD45R/B220** | PE-Cy7 | Rat IgG2a, ϰ | RA3-6B2 | 103222 | Biolegend |
| **CD19** | Alexa Flour 700 | Rat IgG2a K | eBio1D3 | 56-0193-80 | eBioscience |
| **CD11b** | FITC | Rat IgG2b K | M1/70 | 557396 | BD Pharmingen |
| **CD11c** | FITC | Rat IgG2b K | N418 | 117306 | Biolegend |
| **Gr-1** | APC | Rat IgG2b K | RB6-8C5 | 108412 | Biolegend |
| **CD3** | APC | Rat IgG2b K | 145-2C11 | 553066 | BD Pharmingen |
| **CD11b** | APC | Rat IgG2b K | M1/70 | 101212 | Biolegend |
| **Ter 119** | APC | Rat IgG2b K | TER-119 | 116212 | Biolegend |
| **CD19** | APC | Rat IgG2b K | MB19-1 | 17-0191-81 | eBioscience |
| **IL-7Ra/CD127** | Alexa Flour 647 | Rat IgG2b K | A7R34 | 135011 | Biolegend |
| **CD117/cKit** | FITC | Rat IgG2b K | 2B8 | 105805 | Biolegend |
| **CD135/flk2** | PE | Rat IgG2b K | A2F10 | 135305 | Biolegend |
| **CD16/32 or FCRIII/II** | PE-Cy7 | Rat IgG2b K | 93 | 101317 | Biolegend |
| **CD34** | Alexa Fluor 700 | Rat IgG2b K | RAM35 | 56-0341-82 | Invitrogen |
| **Sca-1** | PerCp-Cy5.5 | Rat IgG2b K | D7 | 108123 | Biolegend |
| **CD150** | BV421 | Rat IgG2b K | TC15-12F12.2 | 115925 | Biolegend |
| **CD48** | BV605 | Rat IgG2b K | HM48-1 | 103441 | Biolegend |
| **CD4** | FITC | Rat IgG2b K | RM4.5 | 11-0042-85 | eBioscience |
| **CD44** | PE-Cy7 | Rat IgG2b K | IM7 | 25-0441-81 | eBioscience |
| **CD25** | PerCPCy5.5 | Rat IgG2b K | PC61 | 102029 | eBioscience |
| **CD117 (cKit)** | APC | Rat IgG2b K | 1B8 | 553356 | BD Pharmingen |
| **CD19** | PE-Cy7 | Rat IgG2b K | 6D5 | 115520 | Biolegend |
| **NK1.1** | PerCPCy5.5 | Rat IgG2b K | PK136 | 61-5941-80 | eBioscience |
| **IgM** | APC | Rat IgG2b K | II/4I | 17579082 | eBioscience |
| **CD21/35** | PerCPCy5.5 | Rat IgG2b K | 7E9 | 123415 | Biolegend |
| **CD23** | Alexa Flour 700 | Rat IgG2b K | B3B4 | 101631 | Biolegend |
| **CD45.2** | Pacific Blue | MOUSE SJL | 104 | 109820 | Biolegend |
| **CD45.1** | BV786 | Mouse | A20 | 740889 | BD Biosciences |
| **CD8** | PerCp-Cy5.5 | Rat IgG2a K | 53-6.7 | 553036 | BD Pharmingen |
| **AffiniPure F(ab9)2 Frag goat-anti-human IgG, Fcγ Frag Spec** | APC | Goat-anti-human IgG | | 109-136-170 | Jackson ImmunoResearch |

| | | | | | |
|---|---|---|---|---|---|
| **AffiniPure F(ab9)2 Frag goat-anti-human IgG, Fcγ Frag Spec** | DyLight 405 | Goat-anti-human IgG | | 109-476-170 | Jackson ImmunoResearch |
| **Rhodamine Red-X-conjugated donkey anti-sheep IgG** | Rhodamine Red-X | Donkey anti-sheep IgG | | 713-295-003 | Jackson ImmunoResearch |
| **CD16/ CD32 (mouse Fc block)** | | Rat anti-mouse | 2.4G2 | 553142 | BD Pharmingen |
| **NOTCH1** | | Ag-purified polyclonal sheep | aa19-526 | AF5267 | R&D Systems |
